# Convolutional neural network models of V1 responses to complex patterns

**DOI:** 10.1101/296301

**Authors:** Yimeng Zhang, Tai Sing Lee, Ming Li, Fang Liu, Shiming Tang

## Abstract

In this study, we evaluated the convolutional neural network (CNN) method for modeling V1 neurons of awake macaque monkeys in response to a large set of complex pattern stimuli. CNN models outperformed all the other baseline models, such as Gabor-based standard models for V1 cells and various variants of generalized linear models. We then systematically dissected different components of the CNN and found two key factors that made CNNs outperform other models: thresholding nonlinearity and convolution. In addition, we fitted our data using a pre-trained deep CNN via transfer learning. The deep CNN’s higher layers, which encode more complex patterns, outperformed lower ones, and this result was consistent with our earlier work on the complexity of V1 neural code. Our study systematically evaluates the relative merits of different CNN components in the context of V1 neuron modeling.

## 1 Introduction

There has been great interest in the primary visual cortex (V1) since pioneering studies decades ago (Hubel and Wiesel, 1968, 1959, 1962). V1 neurons are traditionally classified as simple and complex cells, which are modeled by linear-nonlinear (LN) models (Heeger, 1992) and energy models (Adelson and Bergen, 1985), respectively. However, a considerable gap between the standard theory of V1 neurons and reality has been demonstrated repeatedly, at least from two aspects. First, although standard models explain neural responses to simple stimuli such as gratings well, they cannot explain satisfactorily neural responses to more complex stimuli, such as natural images and complex shapes (David and Gallant, 2005; Victor et al., 2006; Hegdé and Van Essen, 2007; Köster and Olshausen, 2013). Second, more sophisticated analysis techniques have revealed richer structures in V1 neurons than those dictated by standard models (Rust et al., 2005; Carandini et al., 2005). As an additional yet novel demonstration of this gap, using large-scale calcium imaging techniques, we (Li et al., 2017; Tang et al., 2018) have recently discovered that a large percentage of neurons in the superficial layers of V1 of awake macaque monkeys respond strongly to highly specific complex features; this finding suggests that some V1 neurons act as complex pattern detectors rather than Gabor-based edge detectors as dictated by classical studies (Jones and Palmer, 1987a; Dayan and Abbott, 2001).

While our previous work (Tang et al., 2018) has shown the existence of complex pattern detector neurons in V1, a quantitative understanding of the relationship between input stimuli and neural responses for those neurons has been lacking. One way to better understand these neurons quantitatively is to build computational models that predict their responses given input stimuli (Wu et al., 2006). If we can find a model that accurately predicts neural responses to (testing) stimuli not used during training, a careful analysis of that model should give us insights into the computational mechanisms of the modeled neuron(s). For example, we can directly examine different components of the model (McIntosh et al., 2017; McFarland et al., 2013; Prenger et al., 2004), find stimuli that maximize the model output (Kindel et al., 2017; Olah et al., 2017), and decompose model parameters into simpler, interpretable parts (Rowekamp and Sharpee, 2017; Park et al., 2013).

A large number of methods have been applied to model V1 neural responses, such as ordinary least squares (Theunissen et al., 2001; David and Gallant, 2005), spike-triggered average (Theunissen et al., 2001), spike-triggered covariance (Touryan et al., 2005; Rust et al., 2005), generalized linear models (GLMs) (Kelly et al., 2010; Pillow et al., 2008), nested GLMs (McFarland et al., 2013), subunit models (Vintch et al., 2015), and artificial neural networks (Prenger et al., 2004). Compared to more classical methods, convolutional neural networks (CNNs) have recently been found to be more effective for modeling retinal neurons (Kindel et al., 2017) and V1 neurons in two studies concurrent to ours (McIntosh et al., 2017; Cadena et al., 2017). In addition, CNNs have been used for explaining inferotemporal cortex and some other areas (Yamins et al., 2013; Kriegeskorte, 2015; Yamins and DiCarlo, 2016). Nevertheless, existing studies mostly treat the CNN as a black box without analyzing much the reasons underlying its success relative to other models, and we are trying to fill that knowledge gap explicitly in this study.

To understand the CNN’s success better, we first evaluated the performance of CNN models, Gabor-based standard models for simple and complex cells, and various variants of GLMs on modeling V1 neurons of awake macaque monkeys in response to a large set of complex pattern stimuli (Tang et al., 2018). We found that CNN models outperformed all the other models, especially for neurons that acted more like complex pattern detectors than Gabor-based edge detectors. We then systematically explored different variants of CNN models in terms of their nonlinear structural components, and found that thresholding nonlinearity and max pooling, especially the former, were important for the CNN’s performance. We also found that convolution (spatially shifted filters with shared weights) in the CNN was effective for increasing model performance. Finally, we used a pre-trained deep CNN (Simonyan and Zisserman, 2014) to model our neurons via transfer learning (Cadena et al., 2017), and found that the deep CNN’s higher layers, which encode more complex patterns, outperformed lower ones; the result was consistent with our earlier work (Tang et al., 2018) on the complexity of V1 neural code. While some of our observations have been stated in alternative forms in the literature, we believe that this is the first study that systematically evaluates the relative merits of different CNN components in the context of V1 neuron modeling.

## 2 Stimuli and neural recordings

### 2.1 Stimuli

Using two-photon calcium imaging techniques, we collected neural population data in response to a large set of complex artificial “pattern” stimuli. The “pattern” stimulus set contains 9500 binary (black and white) images of about 90 px by 90 px from five major categories: orientation stimuli (OT; bars and gratings), curvature stimuli (CV; curves, solid disks, and concentric rings), corner stimuli (CN; line or solid corners), cross stimuli (CX; lines crossing one another), and composition stimuli (CO; patterns created by combining multiple elements from the first four categories). The last four categories are also collectively called non-orientation stimuli (nonOT). See Figure 1 for some example stimuli. In this study, the central 40 px by 40 px parts of the stimuli were used as model input as 40 pixels translated to 1.33 degrees in visual angle for our experiments and all recorded neurons had classical receptive fields of diameters well below one degree in visual angle around the stimulus center (Tang et al., 2018). The cropped stimuli were further downsampled to 20 px by 20 px for computational efficiency. Later, we use ***x**_t_* to represent the t-th stimulus as a 20 by 20 matrix, with 0 for background and 1 for foreground (there can be intermediate values due to downsampling), and 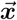 to denote the vectorized version of ***x**_t_* as a 400-dimensional vector.

**Fig. 1.**
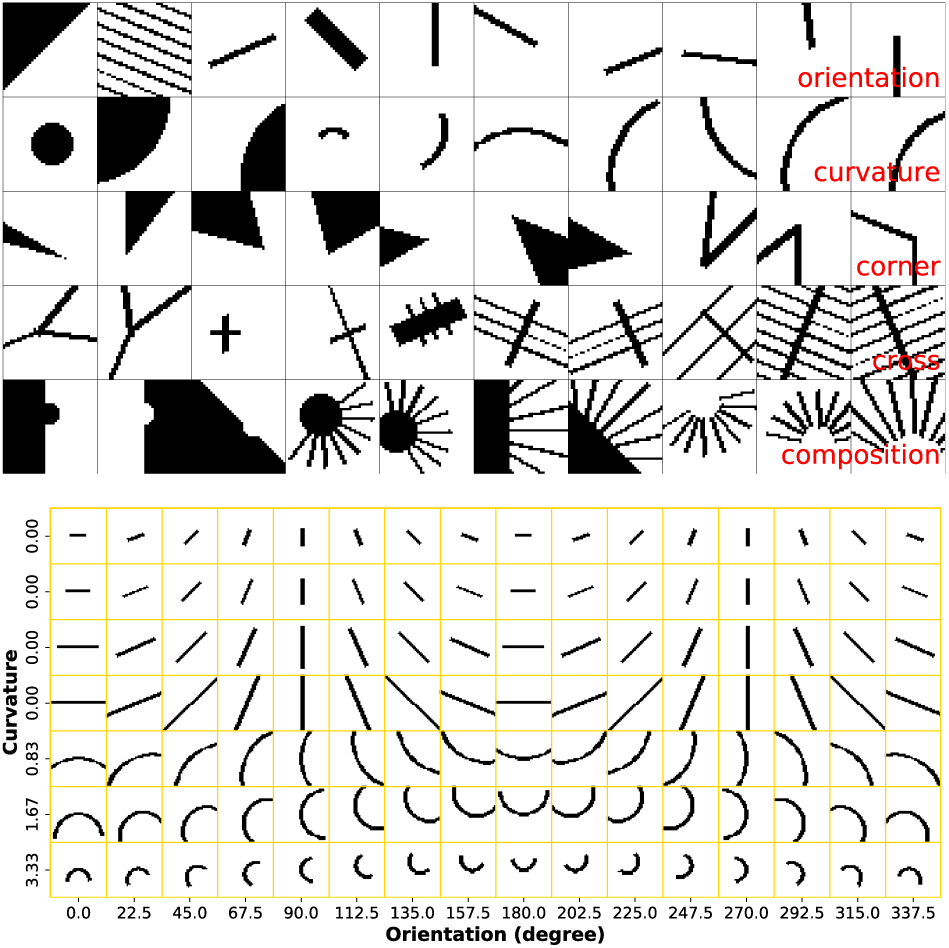
**Top** “Pattern” stimulus set. Stimuli are arranged in rows, each row showing 10 randomly drawn stimuli for each of the five categories (see the bottom right corner of each row). Only the central 40 px by 40 px parts of stimuli are shown. Refer to Tang et al. (2018) for details. **Bottom** A subset of curvature and line stimuli in the stimulus set, ordered by stimulus parameters (curvature, length, and orientation). Only the central 40 px by 40 px parts are shown.

### Stimulus type

Previous work modeling V1 neurons mostly used natural images or natural movies (Kindel et al., 2017; Cadena et al., 2017; David and Gallant, 2005), while we used artificial pattern images (Tang et al., 2018). While neural responses to natural stimuli arguably reflect neurons’ true nature better, it has the following problems in our current study: 1) public data sets (Coen-Cagli et al., 2015) of V1 neurons typically have much fewer images and neurons than our data set, and limited data may introduce bias on the results; 2) artificially generated images can be easily classified and parameterized, and this convenience allows us to classify neurons and compare models over different neuron classes separately (Section 2.2). While white noise stimuli (Rust et al., 2005; McIntosh et al., 2017) are another option, we empirically found that white noise stimuli (when limited) would not be feasible for finding the correct model parameters (assuming CNN models are correct); see Supplementary Materials.

### 2.2 Neural recordings

The neural data were collected from V1 superficial layers 2 and 3 of two macaque monkeys A and B. For monkey A, responses of 1142 neurons in response to all 9500 (1600 OT and 7900 nonOT) stimuli were collected. For monkey B, responses of 979 neurons in response to a subset of 4605 (800 OT and 3805 nonOT) stimuli were collected due to time constraints. Each stimulus was presented for 5 repetitions for both monkeys. During each repetition, all recorded neurons’ responses in terms of *ΔF/F* were collected. Later, we use 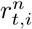 to denote the neural response of the *n*-th neuron for the *t*-th stimulus in the i-th trial (*i* = 1,…, 5), 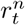 to denote the average neural response over trials, and 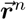 to denote all the average neural responses for this neuron as a vector. Specifically, we have *n* = 1,…, 1142, *t* = 1,…, 9500 for monkey A and *n* = 1,…, 979, *t* = 1,…, 4605 for monkey B.

#### Cell classification

The recorded neurons in the neural data had mixed tuning properties (Tang et al., 2018): some acted more like complex pattern detectors, some acted more like simple oriented edge detectors, and some had weak responses to all the presented stimuli. To allow cleaner and more interpretable model comparisons, we evaluated model performance for different types of neurons separately (Section 5). For example, when comparing a CNN model and a GLM, we computed their performance metrics over neurons that were like complex pattern detectors and those more like simple edge detectors separately, as it is possible that neurons of different types are modeled best by different model classes. To make such per-neuron-type comparison possible, a classification of neurons is required. Here we use the neuron classification scheme in Tang et al. (2018). First, neurons whose maximum mean responses were not above 0.5 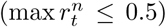 were discarded as their responses were too weak and might be unreliable; then, among all the remaining neurons that passed the reliability test, neurons whose maximum mean responses over nonOT stimuli were more than twice of those over OT stimuli 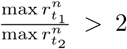 where *t*_1_ and *t*_2_ go over all nonOT and OT stimuli respectively) were classified as HO (higher-order) neurons and the others were classified (conservatively) as OT neurons; finally, all the HO and OT neurons were further classified into subtypes, such as curvature neurons and corner neurons, based on ratio tests similar to the one above—for example, an HO neuron was additionally considered as a curvature neuron if its maximum response over curvature stimuli was more than twice of that over noncurvaturestimuli. Overall, ignoring the unreliable ones, at the top level, there were OT neurons and HO neurons; OT neurons were further classified as classical and end-stopping (neurons that responded well to short bars but poorly to long ones) neurons; HO neurons were further classified as curvature, corner, cross, composition, and mixed (neurons that failed ratio tests for all the four types of nonOT stimuli) neurons. Figure 7 shows example neurons of different classes.

#### Recording technique

While most other studies use spiking data collected using multi-electrode array (MEA) technologies, we use calcium imaging data (Li et al., 2017; Tang et al., 2018). Although MEA-based spiking data are in theory more accurate, calcium imaging techniques can record many more neurons and do not suffer from spike sorting errors. In addition, Li et al. (2017) have shown that the calcium imaging technique we used exhibits linear behavior with MEA technologies across a wide range of spiking activities.

## 3 Methods

Here, we describe three classes of models for modeling V1 neurons in our data set. All the models explored in this study can be considered variants of one-hidden-layer neural networks with different constraints and components. By considering them in the framework of one-hidden-layer neural networks (Section 3.4), we can easily identify key components that make CNNs perform better. In addition, all the methods here model each neuron separately (no parameter sharing among models fitted to different neurons) and the numbers of parameters of different models are kept to be roughly the same if possible; the parameter separation and equality in model size ensure a fairer comparison among models. For each neuron *n* from some monkey, all our models take image ***x**_t_* of size 20 by 20 as input and try to predict the neuron’s mean response 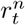 of image t as output. See Section 2 for an explanation of the notation.

### 3.1 CNN models

A CNN model passes the input image through a series of linear-nonlinear (LN) operations—each of which consists of convolution, ReLU nonlinearity (Krizhevsky et al., 2012), and (optionally) max pooling. Finally, outputs of the final LN operation are linearly combined as the predicted response of the neuron being modeled. Our baseline CNN model for V1 neurons is shown in Figure 2, with one (convolutional) layer and 9 filters. Given a 20 by 20 input, it first convolves and rectifies (“convolve + threshold” in the figure) the input with 9 filters of size 9, yielding 9 feature maps (channels) of size 12 by 12, one for each filter. Then max pooling operation (“max pool” in the figure) is performed for each feature map separately to produce 9 pooled feature maps of size 4 by 4. Finally, all the individual output units across all the pooled feature maps are linearly combined (“linear combination” in the figure), plus some bias, to generate the predicted neural response.

**Fig. 2.**
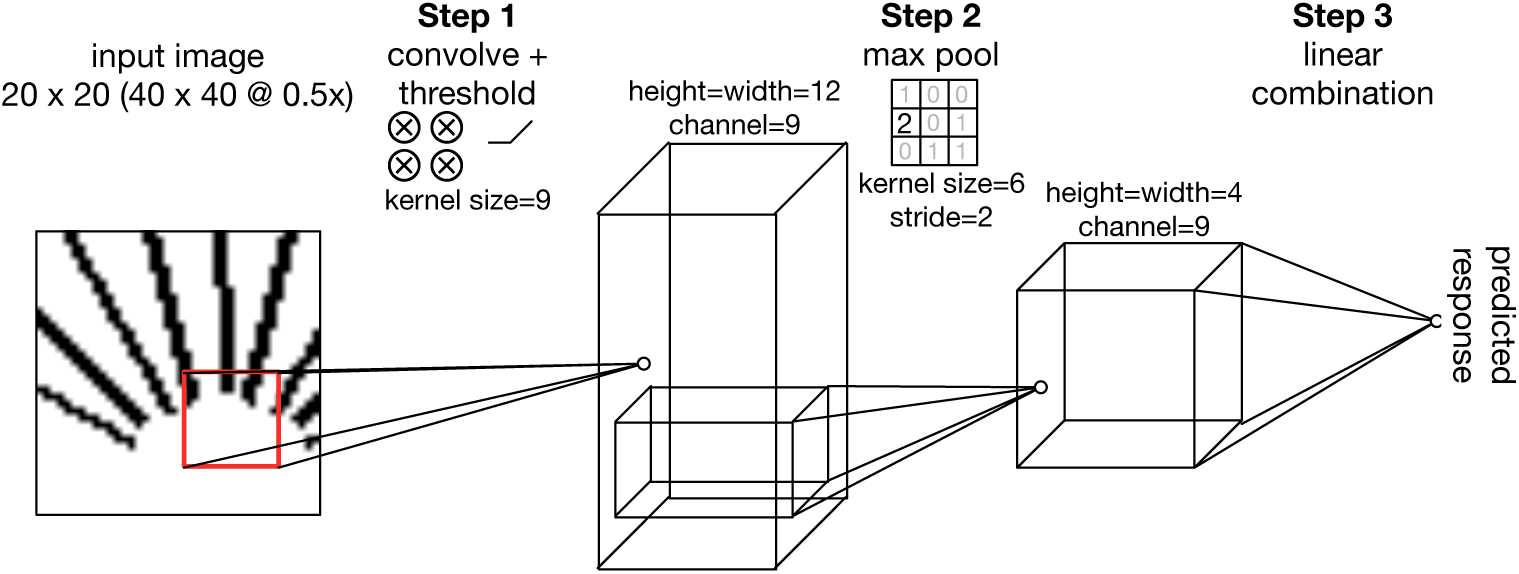
The architecture of our baseline CNN model (or B.9 in Table 2). Given a 20 by 20 input image (40 by 40 downsampled to half; see Section 2), the model computes the predicted neural response in three steps. In **Step 1** (“convolve + threshold”), the model convolves and rectifies the input to generate an intermediate output of size 12 by 12 by 9 (height, width, channel; 3D block in the middle); concretely, for each of the model’s 9 filters of size 9 by 9 (kernel size), the model computes the dot product (with some bias) between the filter and every 9 by 9 subregion in the input (red square being one example), rectifies (*x* ↦ max(0,*x*)) all the dot products, and arranges the rectified results as a 12 by 12 feature map; the process is repeated for each of the 9 filters (channels) and all the 9 feature maps are stacked to generate the 12 by 12 by 9 intermediate output. In **Step 2** ( “max pool”), max pooling operation is performed for each feature map separately to produce 9 pooled feature maps of size 4 by 4; concretely, for each of the 12 feature maps obtained in Step 1, maximum values over 6 by 6 subregions are computed every 2 data points (stride) and arranged as a 4 by 4 pooled feature map; the process is repeated for each of the 9 feature maps to generate the 4 by 4 by 9 pooled output. In **Step 3** (“linear combination”), all the individual output units across all the pooled feature maps are linearly combined plus some bias to generate the predicted neural response. See Section 3.1 as well.

As shown in Table 2 of Section 4.1, apart from the baseline model with 9 channels (B.9 in the table), we also explored other CNN models with the same overall architecture but different numbers of channels.

### 3.2 “Standard” Gabor-based models

Gabor filters are widely used in theoretical models of V1 neurons (Dayan and Abbott, 2001; Jones and Palmer, 1987a; Daugman, 1985). Therefore, we tried to fit (relatively speaking) standard Gabor-based V1 models to our data as control. We tried Gabor simple cell models, Gabor complex cell models, as well as their linear combinations (Figure 3). Interestingly, to the best of our knowledge, such models were not examined in the existing V1 data fitting literature in terms of their performance compared to more popular ones such as GLMs.

**Fig. 3.**
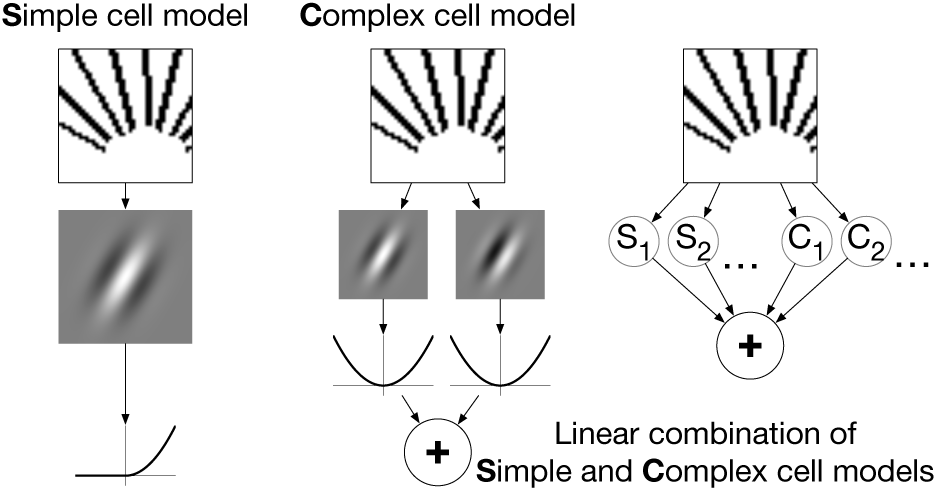
The architecture of Gabor-based models. A simple cell model (left) takes the dot product of a Gabor filter and the input, and passes through the output through a half-wave squaring nonlinearity. A complex cell model (middle) takes the dot products of two Gabor filters with quadrature phase relationship, squares and sums the outputs. A linear combination of simple and complex cell models (right) takes some linear combination of some simple cell models (S) and some complex cell models (C). This figure is partially inspired by Figure 1 in Rust et al. (2005) and Figure 1 in Carandini et al. (2005).

#### 3.2.1 Gabor simple cell models

A Gabor simple cell model (Heeger, 1992; Rust et al., 2005; Carandini et al., 2005) takes the following form:

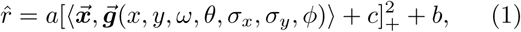

where 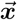 is the input stimulus in raw pixel space re-shaped as a vector (Section 2.1), 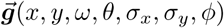 is the Gabor filter (reshaped as a vector) in the raw pixel space with locations *x, y*, frequency *ω*, orientation *θ*, phase *φ*, and standard deviations of the Gaussian envelope *σ_x_, σ_y_*, and *a, b, c* are scale and bias parameters. In such formulation, given some input, the model computes the dot product between the input and its Gabor filter (plus some bias), and passes the output through a half-wave squaring nonlinearity 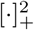. In the existing literature, some simple cell models use threshold nonlinearity (also called over-rectification) [• + *c*]_+_ while some others use half-wave squaring nonlinearity 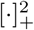; according to Heeger (1992), these two nonlinearities are approximately the same in some sense and therefore we include them both in our models for more flexibility.

#### 3.2.2 Gabor complex cell models

A Gabor complex cell model (Heeger, 1992; Adelson and Bergen, 1985; Rust et al., 2005; Carandini et al., 2005) takes the following form:

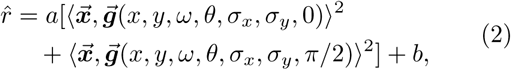

where 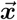 is defined as before, 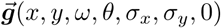 and 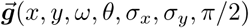 are a pair of Gabor filters (re-shaped as vectors) in the raw pixel space with the same parameters (check Section 3.2.1 for details) except for phases differing by π/2, and *a, b* are scale and bias parameters. In such formulation, given some input, the model computes the outputs of two linear Gabor filters with quadrature phase relationship and sums their squares together to achieve phase invariance. As an aside, while we set phases of the Gabor filter pair to be *φ* and *φ* + *π*/2 with *φ* = 0, any other *φ* will also work; empirically we found Eq. (2) has no or little dependence on *φ*.

#### 3.2.3 Linear combinations of complex and simple cell models

While simple and complex cell models are the canonical ones in most neuroscience textbooks, detailed analyses on monkey V1 neurons have revealed more than one (simple) or two (complex) linear components (Rust et al., 2005; Carandini et al., 2005). Rust et al. (2005) call such extensions to “standard” simple and complex cell models “generalized LNP response models” (Figure 1C of Rust et al. (2005)). One simple realization of generalized LNP response models is to take linear combinations of “standard” Gabor-based models that are defined above.

### 3.3 Generalized linear models

We consider the following set of Poisson generalized linear models (McCullagh and Nelder, 1989; Paninski, 2004) with possibly nonlinear input transformations (Figure 4). We also tried Gaussian GLMs and they performed consistently worse than Poisson ones in our experiments. Note that the term “GLM” has been used pretty loosely in the literature, and many models with similar structural components to those in the CNN are considered GLMs by many. We want to emphasize that the purpose of including GLMs in this study is not to compare CNNs and (all the variations of) GLMs in terms of performance but to find key components that make CNN models outperform commonly used models for V1 modeling. We call these models GLMs mainly because they are often formulated as GLMs in the literature. See Section 3.4 for the connection between CNNs and GLMs considered in this study.

**Fig. 4.**
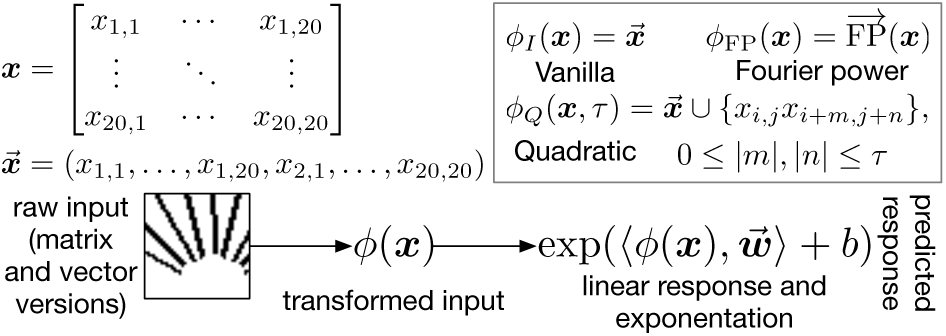
The architecture of generalized linear models. The raw input stimulus *x* is first transformed into *φ(x),* where different *φ*(·) are used for different GLM variants (inside the box). For vanilla GLMs, we use the identity transformation *φ_I_*(·); for Fourier power models, we use the Fourier power transformation *φ*_FP_(·) (Section 3.3.2); for generalized quadratic models, we use the localized quadratic transformation *φ_Q_*(*·,τ*) (Section 4.3.2 and Section 3.3.3). The transformed input *φ*(***x***) is passed into a linear function ⟨·,**w**⟩ + *b* and the output is exponentiated to give the predicted neural response. For details on the localized quadratic transformation (*φ*_*Q*_(·,*τ*) in the figure), see Section 4.3.2 and Figure 5.

#### 3.3.1 Vanilla generalized linear models

A vanilla (linear) GLM takes the following form:

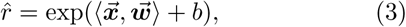

where 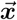 is the input stimulus in raw pixel space re-shaped as a vector, 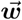 is the linear spatial filter in the raw pixel space, and *b* is the bias parameter for adjusting the firing threshold. The formulation of this vanilla GLM is standard for modeling V1 simple cells (Jones and Palmer, 1987b; David and Gallant, 2005; Carandini et al., 2005), which respond to stimuli having appropriate orientation, spatial frequency, and spatial phase (Hubel and Wiesel, 1959).

#### 3.3.2 Fourier power models

A Fourier power model (David and Gallant, 2005) takes the following form:

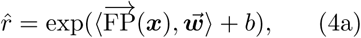

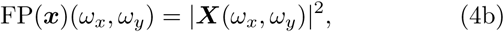

where FP(x) computes the Fourier power spectrum of the input stimulus (FP(***x***) is 2D in Eq. (4b) and reshaped to a vector 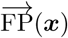 in Eq. (4a)), ***X*** denotes the 2D Fourier transform of the input stimulus, 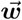 is the linear filter in the Fourier power domain, and *b* is the bias parameter for adjusting the firing threshold. In practice, Fourier power models provide performance close to the state of the art (Carandini et al., 2005; Köster and Olshausen, 2013).

#### 3.3.3 Generalized quadratic models

A generalized quadratic model (GQM) (Park and Pillow, 2011; Park et al., 2013) takes the following form:

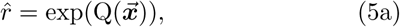

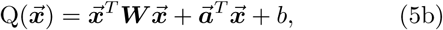

where 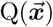 computes a quadratic feature transformation of the (vectorized) input stimulus, ***W***, 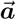, *b* are the second-order parameters, first order parameters, and bias parameter respectively in the transformation. A GQM can be formulated as a GLM with quadratic feature transformation (Park et al., 2013), which introduces additional nonlinearity components and flexibility for neuron modeling. In addition, there is a connection between GQMs and spike-triggered based methods under certain conditions (Park and Pillow, 2011) and GQMs are statistically more efficient. Note that Fourier power models (Section 3.3.2) can be also formulated as GQMs, as 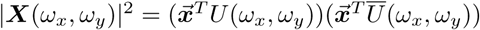 in Eq. (4b), where *U*(*ω_x_*, *ω_y_*) denotes the Fourier transform vector for frequency pair (*ω_x_*, *ω_y_*) and 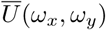 denotes its complex conjugate.

### 3.4 Connections among CNNs, Gabor models, and GLMs

As mentioned in the beginning of Section 3, the three classes of models considered in this study are connected and form a continuum as they all can be roughly formulated as vanilla one-hidden-layer neural networks (Bishop, 2006), or one-hidden-layer multilayer perceptrons (MLPs):

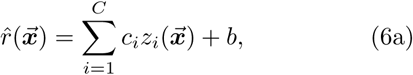

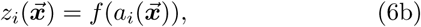

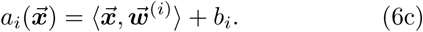

A one-hidden-layer neural network computes the output 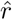 given (vectorized) input stimulus 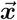 according to Eqs. (6). Overall, the output is a linear combination of *C* hidden units’ output values *z_i_* as shown in Eq. (6a). Each hidden unit’s output is computed by applying some nonlinearity (also called activation function) *f* on the pre-activation value of the hidden unit *a_i_* as shown in Eq. (6b), and pre-activation value *a_i_* is a linear function of input specified by weights 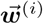 and bias *b_i_* as shown in Eq. (6c).

Gabor models can be formulated as MLPs with constraints that weights 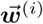 must be Gabor functions. A simple cell model is a MLP with one hidden unit and half-wave squaring nonlinearity; a complex cell model is a MLP with two hidden units in quadrature phase relationship and squaring nonlinearity; a linear combination of simple and complex cell models is a MLP with multiple hidden units and mixed nonlinearities.

GLMs can be formulated as MLPs with an additional exponential nonlinearity on output. A vanilla GLM is a MLP with one hidden unit and no nonlinearity (linear); a Fourier power GLM is a MLP with multiple hidden units of fixed weights (Fourier basis functions) and squaring nonlinearity; A GQM is a MLP with multiple hidden units and squaring nonlinearity— the linear term in Eq. (5b) can be absorbed into the quadratic one as long as the quadraic coefficient matrix is full rank. Empirically, we found the additional accelerating exponential nonlinearity to be unimportant for the modeling of our data, as Poisson GLMs with the additional accelerating nonlinearity performed similarly or marginally better, compared to Gaussian and softplus GLMs without such nonlinearity (Supplementary Materials).

A CNN can be formulated as a MLP with ReLU (*x* ↦ max(0, *x*)) nonlinearity and an additional max pooling operation before the final output computation of Eq. (6b). Compared to other models, a CNN has additional constraints among the weights of hidden units— shared and spatially shifted in groups. For example, our baseline CNN can be considered as a MLP with 12 × 12 × 9 = 1296 hidden units, as each 9 by 9 filter in the CNN yields a feature map of 12 × 12 = 144 hidden units, and there are 9 filters in the CNN. For MLP hidden units derived from a common feature map, filter weights are shared and spatially shifted; for MLP hidden units derived from different feature maps, filter weights are independent. This group-wise sharing of hidden unit weights in CNN models is not present in GLMs, which we will compare in detail with CNNs in Section 5 as GLMs were the best-performing non-CNN models in our experiments.

Table 1 gives a summary of different models in terms of their structures, under the framework of one-hidden-layer neural network (or MLP). We classify nonlinearities into thresholding (half-wave squaring and ReLU) and non-thresholding (squaring) ones, because we found all the thresholding activation functions behaved essentially the same in our experiments (Section 5.2) and we think that being thresholding or not may be the most important aspect for a nonlinearity.

**Table 1.** Comparison of model structures for Gabor models, GLMs, and CNNs in the framework of one-hidden-layer MLP. First two columns specify the model class and subclass. The third column shows whether the models’ corresponding MLPs have multiple hidden units or not. The fourth column shows the constraints among hidden units imposed by the models; “independent” means weights for different hidden units can vary independently, “shared” means weights for different hidden units are tied together (via convolution), “quadrature phase” means weights of the hidden unit pair are in quadrature phase relationship (specific to Gabor models), and “fixed” means weights are not learned but specified before training. The fifth column specifies the nonlinearity (activation function), with “none” meaning no nonlinearity (identity or linear activation function), and “mixed” meaning both thresholding and non-thresholding nonlinearities. The last column specifies additional structures imposed by the models.

| Class | Subclass | Multiple units | constraints among units | nonlinearity | additional structures |
| --- | --- | --- | --- | --- | --- |
| Gabor | simple | No | — | thresholding | weights are Gabor |
|  | complex | Yes | quadrature phase | non-thresholding | weights are Gabor |
|  | combination | Yes | independent | mixed | weights are Gabor |
| GLM | vanilla | No | — | none | exponential output |
|  | Fourier power | Yes | fixed (not learned) | non-thresholding | exponential output |
|  | GQM | Yes | independent | non-thresholding | exponential output |
| CNN | — | Yes | independent + shared | thresholding | max pooling |

### 3.5 Pre-trained CNNs

There are at least two different ways to model V1 neurons and neural data in general using CNNs: data-driven and transfer learning. In the data-driven approach, CNNs are trained from scratch to fit the neural data. This is the approach taken in this study and many other very recent ones (Kindel et al., 2017; McIntosh et al., 2017). In the transfer learning (also called goal-driven) approach (Yamins and DiCarlo, 2016; Kriegeskorte, 2015), CNN models are first trained on some other tasks such as image classification, and then neural data are fitted by (linearly) combining outputs of fitted units in the trained models. As shown in Cadena et al. (2017), two approaches work similarly for V1 neurons in response to natural images.

As an additional experiment, we tried to model neurons in our data set using a transfer-learning approach similar to that in Cadena et al. (2017). Specifically, we fed all images^1^ to the CNN model VGG19 (Simonyan and Zisserman, 2014) and extracted intermediate feature representations of the images across all the CNN layers (except fully-connected ones). The intermediate representations of each layer were used as inputs to train (a set of) GLMs to model all the neurons. All the other implementation details were the same as those for GLMs (Section 4.3).

### 3.6 Model evaluation

Given some model 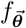 with trainable parameters 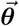, we evaluate its performance 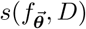 on a single neuron *n* based on its input stimuli and responses 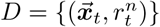, where 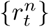 are the across-trial average responses computed from 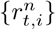 (Section 2.2), by squared normalized correlation coefficient 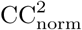 (Schoppe et al., 2016; Hsu et al., 2004).

To evaluate a model, we first partition *D* into training set *D*_train_ and testing set *D*_test_ using 80 % and 20 % of the whole data set, respectively. For those models involving model selection in the training (all but Gabor models), 20 % of the training data is reserved for validation purpose. We use D_*train*_ to obtain the trained model 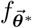, and compute the model performance 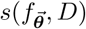 as follows (neuron index *n* is omitted as there is only one neuron being considered):

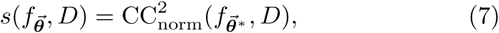

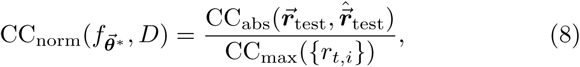

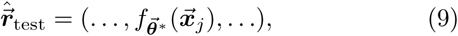

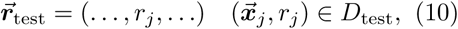

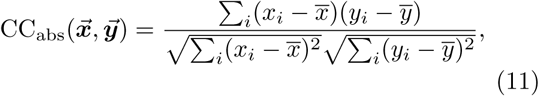

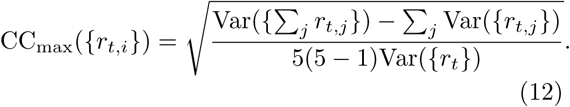

Concretely, we first compute the raw Pearson correlation CC_abs_ between the set of neural responses 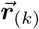 and the set of model responses 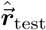 using Eq. (11) (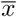 and 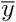 denote mean values of the two inputs), then divide this CC_abs_ by CC_max_, which is defined in Eq. (12) (adapted from Schoppe et al. (2016), with 5 in the denominator being the number of trials) and estimates the maximal Pearson correlation coefficient an ideal model can achieve given the noise in the neural data (Schoppe et al., 2016; Hsu et al., 2004), to get the normalized Pearson correlation coefficient using Eq. (8), and finally square CC_norm_ to get the model performance 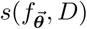 using Eq. (7). As squared CC_abs_ gives the fraction of variance in neural responses explained by the model in a simple linear regression, squared CC_norm_ gives the normalized explained variance that accounts for noise in the neural data. Notice that CC_max_ is computed over all the data instead of testing data for more accurate estimation.

Our definition of model performance depends on how D is partitioned. To make our results less susceptible to the randomness of data partitioning, we report results averaged over two partitions.

## 4 Implementation Details

### 4.1 CNN models

#### 4.1.1 Detailed model architecture

Table 2 shows all the three CNN model architectures we evaluated against other models (Section 5), with the baseline CNN model (Figure 2) denoted B.9 in the table. For a fair comparison between CNNs and other models (primarily GLMs; Gabor models inherently have too few parameters), in addition to the baseline CNN model B.9, we also evaluated two variants of the baseline model by changing its number of channels. Overall, the three CNN models match the three classes of GLMs (Section 4.3) in terms of model size. For vanilla GLMs (401 parameters), we picked the 4-channel CNN architecture (393 parameters); for Fourier power GLMs (around 200 parameters due to the symmetry of Fourier power for real-valued input), we picked the 2-channel one (197 parameters). For GQMs, whose original numbers of parameters are too large, we decided to perform PCA on their input data to reduce the dimensionality. We set the reduced dimensionality to 882 (therefore 883 parameters for GQMs), and evaluated GQMs against the baseline 9-channel CNN architecture (883 parameters). While we could keep more or fewer input dimensions for GQMs and use CNN models with more or fewer channels accordingly, we found that (1) the CNN’s performance relatively plateaued for having more than 9 channels (see Supplementary Materials) and (2) keeping fewer input dimensions could potentially affect the performance of GQMs (see Section 4.3.2). The 2- and 4-channel CNNs were used mainly for a fair comparison of CNNs and other models, and the baseline 9-channel one was further analyzed.

**Table 2.**
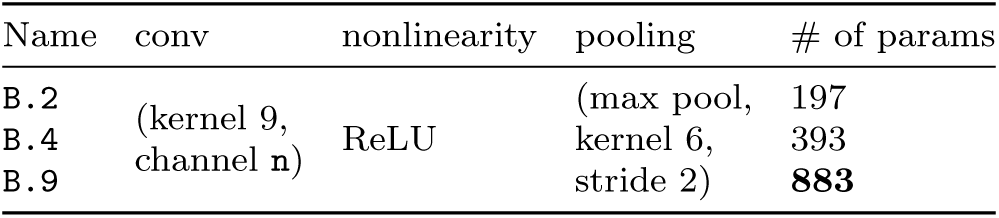
CNN model architectures explored in this work. Each row describes one CNN model architecture, with the first column showing its name (B.n where n is the number of channels), middle columns describing its computational components, and the last showing its number of parameters. Each CNN model first passes the input image through three computational components shown in the table—convolution (conv), nonlinearity, and pooling—and then linearly combine (“fully connected” in CNN jargon) output values of the pooling operation to give the model output. The baseline CNN (B.9) has its number of parameters shown in boldface. The number of parameters is computed by adding the number of parameters in the convolutional layer and that in the fully connected layer. For example, the baseline model B.9 has 9 × (9 × 9 + 1) = 738 parameters (9 for number of channels, 9 for kernel size, and 1 for bias) for the convolutional layer, and 9 × 4 × 4 + 1 = 145 parameters (9 for number of channels, 4 for pooled feature map’s size, and 1 for bias) for the fully connected layer, resulting in 738 + 145 = 883 parameters.

#### 4.1.2 Optimization

The models were implemented in PyTorch (Paszke et al., 2017), version 0.3.1. Model parameters were optimized to minimize the mean squared error between model outputs and recorded neural responses. Of all the training data, 80 % of them were used for actual training, and the remaining 20 % were kept as validation data for early stopping (Goodfellow et al., 2016) and model selection. For each combination of neuron and model architecture, we trained the model four times using four sets of optimization hyperparameters (Table 3), which were selected from more than 10 configurations in our pilot experiments (Supplementary Materials) conducted on about 10 neurons^2^. Of all the four models trained using different optimization hyperparameters, the one with the highest performance on validation data in terms of Pearson correlation coefficient was selected.

**Table 3.** Optimization hyperparameters for CNN models. Minibatch size was set to 128 in all cases, momentum (for SGD) was set to 0.9, and other hyperparameters, such as *β* in Adam (Kingma and Ba, 2014), took default values in PyTorch. LR, learning rate; L2 conv, L2 weight decay on the convolutional layer; L2 fc, L2 weight decay on the fully connected layer; SGD, vanilla stochastic gradient descent with momentum.

| Name | Optimizer | LR | L2 conv | L2 fc |
| --- | --- | --- | --- | --- |
| 1e-3_1e-3_a002 | Adam | 0.002 | 0.001 | 0.001 |
| 1e-4_1e-3_a002 | Adam | 0.002 | 0.0001 | 0.001 |
| 1e-3_1e-3_s1 | SGD | 0.1 | 0.001 | 0.001 |
| 1e-4_1e-3_s1 | SGD | 0.1 | 0.0001 | 0.001 |

### 4.2 “Standard” Gabor-based models

The models were implemented in PyTorch (Paszke et al., 2017). Input stimuli were preprocessed to have zero mean for each stimulus. Model parameters were optimized to minimize the mean squared error between model outputs and recorded neural responses using Adam (Kingma and Ba, 2014) without weight decay and with full batch learning (so gradients were computed using the full data set). To (partially) avoid getting trapped in local optima, for each fitted model, we repeated optimization procedures over hundreds of random initializations and took the set of optimized parameters with the smallest error as the final set of optimized parameters. Empirically, such nested optimization procedure converged to ground-truth model parameters almost all the time. Unlike CNNs (Section 4.1.2) or GLMs (Section 4.3.3), here all the training data were used in the actual training and no model selection was performed, as Gabor models have very few parameters and overfitting should not be a problem.

#### 4.2.1 Linear combinations of “standard” Gabor-based models

The implementation details are essentially the same as those in Section 4.2. We tried the following combinations of simple cell and complex cell models: one simple plus one complex; one simple plus two complex; two simple plus one complex.

### 4.3 Generalized linear models

#### 4.3.1 Vanilla GLMs and Fourier power models

For vanilla GLMs (Section 3.3.1), raw stimuli were vectorized into 400-dimensional vectors as model input; for Fourier power models (Section 3.3.2), we first applied a Hann window to each raw stimulus as done in David and Gallant (2005) to reduce edge artifacts, and then computed 2D Fourier power spectra individually for windowed stimuli as model input.

#### 4.3.2 GQMs

GQMs (Section 3.3.3) are simply standard GLMs with quadratic transformation on input. A full quadratic transformation over a 400-dimensional raw input vector would result in a vector of more than 80,000 dimensions. To make the number of parameters manageable, we performed the following two optimizations for the model input of GQMs.

– Instead of the full quadratic transformation, we performed local quadratic transformations with different “localities” (Figure 5). Local quadratic transformations only compute quadratic terms over stimulus pixels that are close enough. For example, a local transformation with locality 2 will only compute quadratic terms over pixels that are at most 2 pixels apart in both horizontal and vertical axes. For this study, we tried localities 2, 4, and 8.
– Even with local quadratic transformations, the input dimensionality is still too high for efficient op-timization of model parameters. Therefore, we performed principal component analysis (PCA) on the outputs of local quadratic transformations to reduce their dimensionalities. For a fair comparison of GQM models and our CNN models, 882 dimensions ^3^ were kept as this would make our GQMs have the same number of parameters as our 9-channel CNN (see Section 4.1.1; the 9-channel CNN has 883 parameters, and a GLM with a 882-dimensional input has 883 parameters due to the bias parameter). The dimensionality reduction procedure kept over 95 % of the variance; if the input dimensionality had been made to align with CNN models with fewer channels, less than 95 % of the variance would have been kept and the performance of GQMs might have been affected much.

**Fig. 5.**
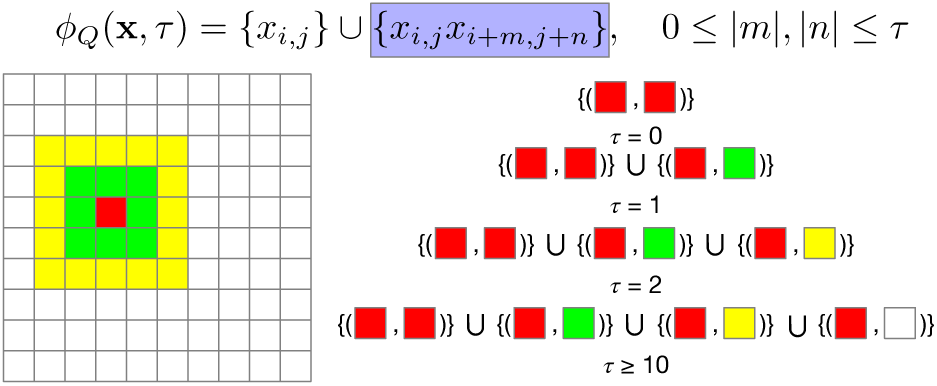
Quadratic feature transformation*φ_Q_*(*x*,*τ*) transforms original stimulus *x*, whose elements are indexed by pixel locations *i, j*, into quadratic features with “locality” *τ*. The output vector contains the union of first order terms {*x_i_,_j_*} and second order terms involving pixels differing by at most *τ* pixels in all directions, as shown by the equation above. The diagram below shows how to compute the second order terms (shaded box in the equation) of some pixel (denoted in red) in a 10 px by 10 px stimulus for different *τ*’s. When *τ* = 0, only the second order interaction between the red pixel and itself is included; when *τ* = 1, additional interactions between the red pixel and each green one are included, and so on.

#### 4.3.3 Optimization

All the models were implemented using glmnet (Friedman et al., 2010) as Poisson GLMs. Similar to CNNs (Section 4.1.2), 80% of the training data were used for actual training and the remaining 20 % were kept as validation data for model selection. L1 regularization was used and the best regularization parameter was selected (out of 100 candidates) by computing model performance on the validation data in terms of Pearson correlation coefficient. For all the three GLM variants, the (transformed) input stimuli have highly correlated dimensions, due to high correlations between adjacent pixels in the original stimuli, and such high correlations in the input made glmnet converge extremely slowly in practice. We worked around this issue by performing full PCA (without reducing the dimensionality) on input stimuli before feeding them to glmnet. Empirically we found this speedup trick made little or no difference to model performance. We also tried Gaussian and softplus GLMs; they performed similarly to or worse than Poisson ones in our experiments (Supplementary Materials).

## 5 Results

### 5.1 CNN models outperformed others especially for higher-order neurons

Figure 6 shows the performance of CNN models vs. others (except pre-trained CNN models; see Section 5.4) on explaining our V1 neural data. Because the full stimulus set consists of different types of stimuli (OT, CN, CV, etc.; see Section 2.1), and the full population of neurons for each monkey consists of two subsets (OT neurons and HO neurons, which can be divided into finer subsets as well; see Section 2.2) that responded very differently to different types of stimuli, we trained all models using different stimulus subsets (“OT” stimuli and all stimuli; we also tried training only on “nonOT” stimuli, and that gave similar results to using all stimuli), and evaluated each model in terms of its average 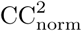 (Section 3.6) averaged over OT neurons and HO neurons (for results on finer subsets, see Section 5.2 and later). We do not show results of HO neurons trained on OT stimuli, as HO neurons by definition did not respond to OT stimuli well and the results might be unreliable.

**Fig. 6.**
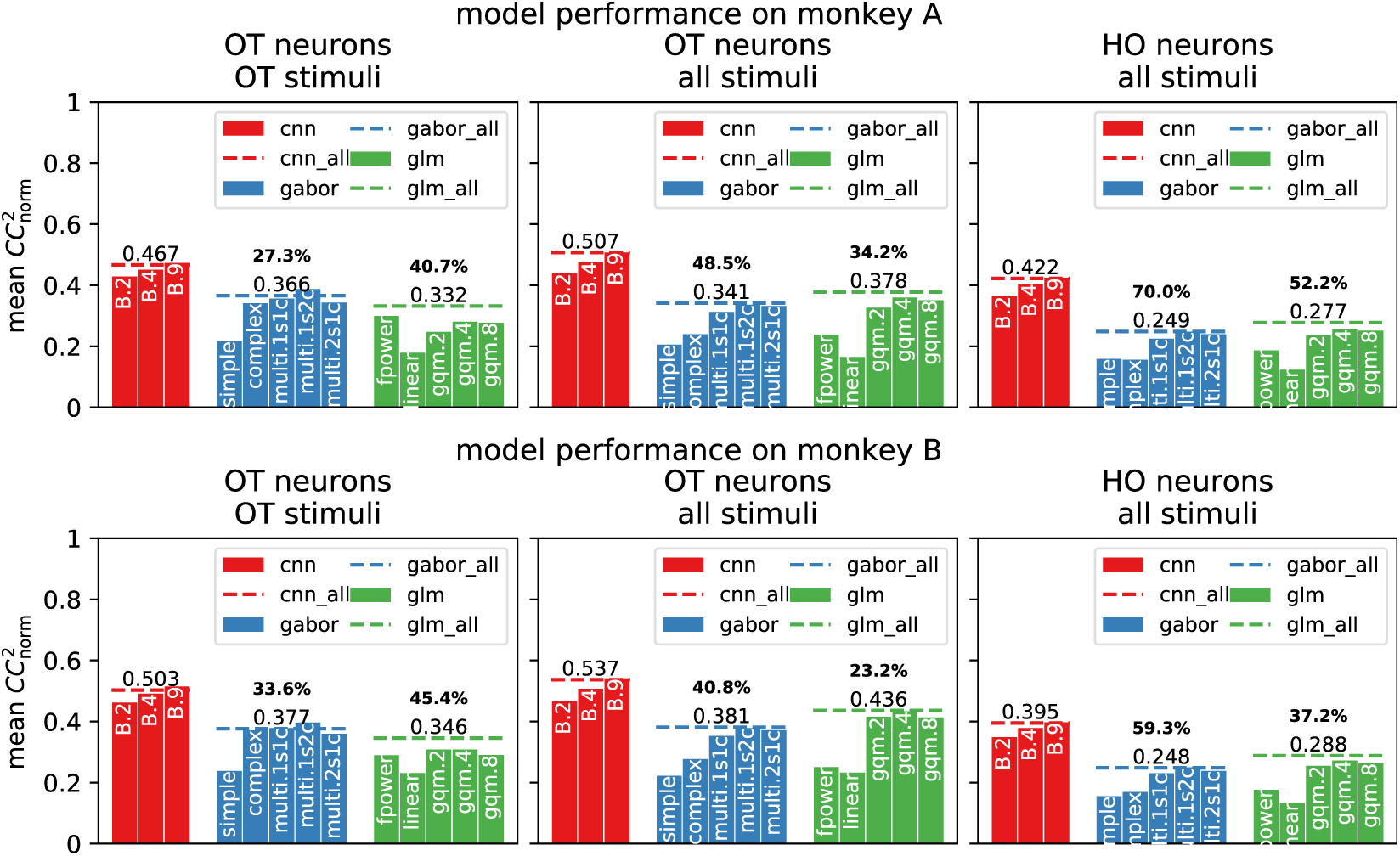
CNN models vs. others on explaining V1 neural data. Two rows of panels show results for monkey A and monkey B respectively, and three columns of panels show how models performed on different neuron subsets (“OT” and “HO”), evaluated on different subsets of stimuli (“OT” and “all”). For each panel, the model performance is shown in 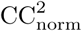 averaged over neurons in the neuron subset. For each category of models (`cnn`, `glm`, etc.), solid bars show model performance of different specific model architectures, and dashed lines (suffixed by `_all`) show the category’s overall “best” performance by taking the best model architecture for each individual neuron (in terms of validation performance for CNNs and GLMs and training performance for Gabor models). Boldface numbers are the relative performance increases of the CNN classes over non-CNN classes (computed as ratios between dashed lines minus one). For CNN models (red), check Table 2 for their meanings. For Gabor models (blue), `complex` and `simple` mean complex cell and simple cell models; `multi.MsNc` means linear combinations of `M` simple and `N` complex model(s). For generalized linear models (green), `linear` means vanilla GLM; `fpower` means Fourier power GLM; `gqm.x` (`x` being one of `0`,`2`,`4`,`8`) means the quadratic GLM with locality `x`.

We compare CNN models and other models at two different levels. At the individual model architecture level (solid bars in Figure 6), we compare specific CNN architectures (models with different numbers of channels) with Gabor models and GLMs. In this case, CNN models with more channels worked better and they outperformed their GLM counterparts (B.2 vs. Fourier power GLMs, B.4 vs. linear GLMs, and B.9 vs. GQMs; see Section 4.1.1) across the board; GQMs had in general better performance than other GLMs, but still fell behind CNNs by a large margin. Gabor models performed similarly to GLMs or worse, and were outperformed by CNNs as well.

At the overall model category level (dashed lines in Figure 6), we compare CNN models as a whole to Gabor models as a whole as well as GLMs as a whole. To do this, for each model category, we constructed an “all” model for that category by choosing the best performing model architecture (in terms of performance on validation data for CNNs and GLMs, and in terms of performance on training data for Gabor models; testing data was never used during the model selection) for each individual neuron. By comparing the dashed lines, we have the following empirical observations about the three model classes.

### CNNs outperformed other models especially for HO neurons with complex stimuli

When stimuli were the same, the relative performance gap between CNN and other models was larger for HO neurons than OT neurons (middle and right columns of panels of Figure 6). For example, on Monkey A, the relative performance increase of the CNN over the GLM increased from 34.2% for OT neurons to 52.2% for HO neurons. When neurons to model were the same, the relative performance gap was larger for complex stimuli than simple stimuli (left and middle columns of panels of Figure 6). For example, on Monkey A, the relative performance increase of the CNN over the Gabor model increased from 27.3% for “OT” stimuli to 48.5 % for all stimuli.

#### Priors on Gabor models helped especially with limited data

When the stimuli were limited and simple, Gabor models outperformed GLMs, possibly due to the strong and neurophysiologically reasonable prior on Gabor models that filter weights can be described well by Gabor functions (Jones and Palmer, 1987a), and vice versa when the stimuli were relatively sufficient and rich (leftmost column of panels vs. other panels of Figure 6). One may hypothesize that multi-component Gabor models (`multi` ones) outperformed standard ones (`complex` and `simple`) mostly due to having multiple orientations; this was not true as shown in Section 5.3.

Finally, Figure 7 shows the fitting results of some neurons in different classes (see Section 2.2); for CNN models, we also show the learned filters and visualization results obtained by activation maximization (Olah et al., 2017); these visualization results are images that activate fitted CNNs most. In most cases, Gabor models and GLMs failed to predict the high-responding parts of the tuning curves compared to CNNs.

**Fig. 7.**
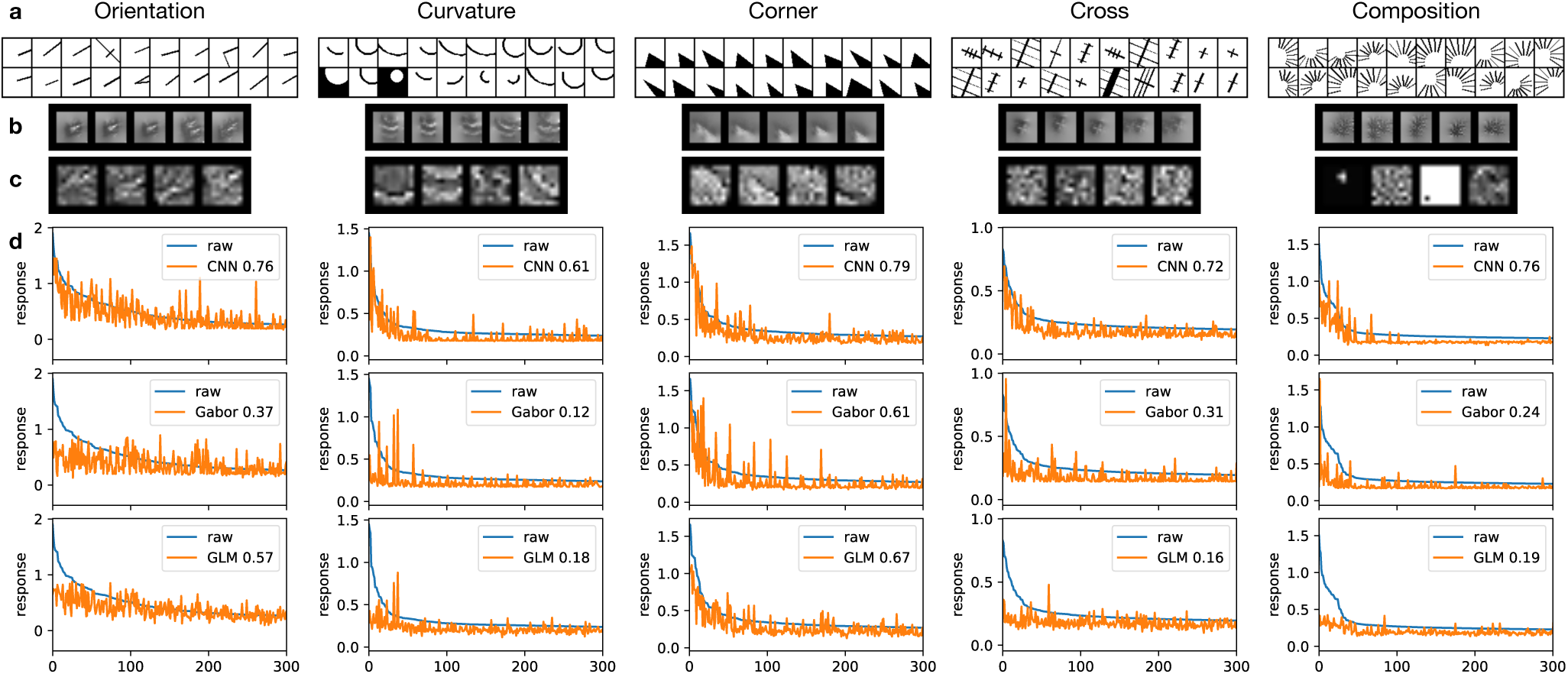
Example neurons and their fitting results. For each of the five stimulus classes shown in different columns, we show the following four pieces of information regarding the fitting of a neuron that responded better to this class than the others (**a-d**). **a** The top 20 responding stimuli of the neuron; **b** the fitted CNN fully connected output layer’s visualization results (over 5 random initalizations) obtained by activation maximization (Olah et al., 2017) implemented in `keras-vis` (Kotikalapudi, 2017); **c** the fitted CNN’s four 9 by 9 convolutional filters (each scaled independently for display); **d** the neuron’s fitting results (over testing data) on three categories of models: CNN, Gabor and GLM, with model performance in terms of 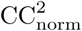 given in the legends. As each category of models has multiple variants or architectures, we roughly speaking picked the overall best one for each category. We picked the 4-channel architecture B.4 for CNN, as it performed almost the same as the baseline B.9 (Figure 6) and allows easier visualization and interpretation; we picked `multi.1s2c` for Gabor, and `gqm.4` for GLM as they performed overall better than other variants. Check Figure 6 for the meanings of model names.

In Supplementary Materials, we show that CNN models outperformed others even with less amount of data; we also show additional results on CNN models, such as comparison of different optimization configurations and comparison of different architectures (different numbers of layers, different kernel sizes, and so on). We will focus on the one-convolutional-layer CNN model B.9 with 883 parameters for the rest of this study, because its performance was close to the best among all the CNN models we tried (Supplementary Materials) without having too many parameters, and its one-layer architecture is easier to analyze than those of similarly performing models.

### 5.2 What made CNNs outperform other models

As shown in Figure 6, the baseline CNN architecture alone (B.9) outperformed GLMs, which were the best non-CNN models in this study, by a large amount, especially for HO neurons. By comparing the row for the CNN and the rows for GLMs (particular the row for the GQM, as GQMs overall performed better than other GLM variants) in Table 1 (Section 3.4), we hypothesize that this performance gap was primarily due to the structural components present in the CNN but not in GLMs we studied: thresholding nonlinearity (ReLU), max pooling, and shared weights of hidden units (convolution). To test our hypothesis, we explored different variants of our baseline CNN architecture B.9 in terms of its structural components. The results on thresholding nonlinearity and max pooling are given in this part, and those on convolution are given in the next part. While our GLMs possess an exponentiation non-linearity which is not present in our CNNs, we found that the exponentiation gave little performance increase than without (Supplementary Materials).

To better understand the utilities of thresholding nonlinearity and max pooling, we explored various variants of the baseline CNN architecture in terms of non-linearity and pooling scheme. Specifically, we tried all combinations of five different nonlinearities—ReLU (**R**), ReLU followed by squaring (half-squaring, **HS**), squaring (**S**), absolute value (**A**), linear (no nonlinearity, **L**)**—** and two different pooling schemes—max pooling (**max**), average (mean) pooling (**avg**)—with other structural components unchanged. Thus, we obtained ten different CNN variants (including the original one) and compared them with the “all” model for GLMs (picking the best model architecture for each neuron), or **GLM_all** as reference. Results are shown in Figure 8 and Figure 9, which have the same organization: panels a-c show the performance of all explored models as before, but with 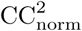 over OT and HO neurons decomposed into average 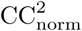 for finer subsets inside OT and HO neurons (Section 2.2) to examine model performance in more detail; panels d-f show the neuron-by-neuron comparison of different pairs of models for highlighting. Overall, we have the following observations (letters in the parentheses denote the panels used for highlighting among d-f, if any).
– Thresholding nonlinearities outperformed non-thresholding ones (d,e).
– Thresholding nonlinearities performed similarly (f).
– No consistently better pooling type, but max pooling was more powerful in isolation.
– High correlation between per-neuron and average model performance (almost all panels).

**Fig. 8.**
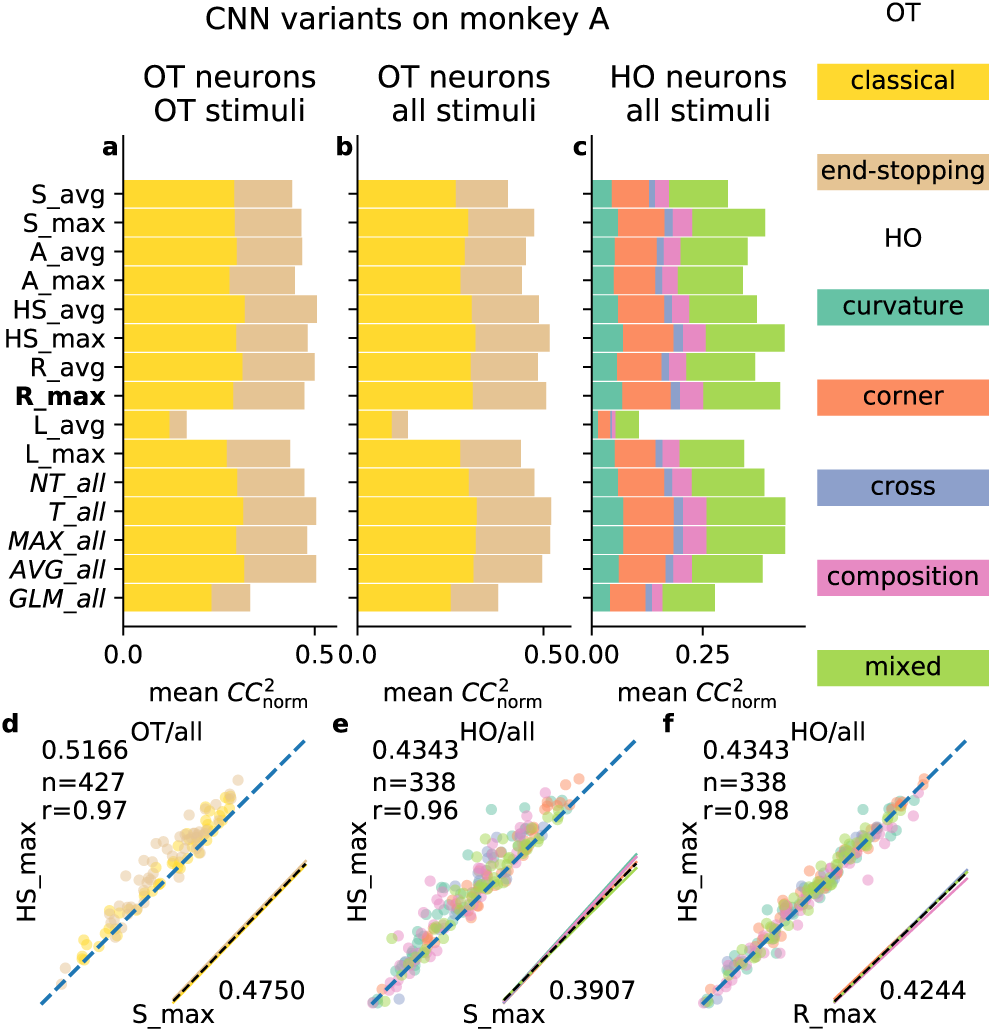
Detailed comparison of CNN variants, monkey A. a-c ten variants of the baseline CNN (B.9), along with the “all” model for GLMs `GLM_all` (Figure 6) for reference. In addition, four “all” CNNs, each of which constructed from CNN models with some shared structural component (thresholding nonlin-earity `T`, non-thresholding nonlinearity `NT`, max pooling `MAX`, or average pooling `AVG`), are shown as well. CNN variants are named `X_Y` where `X` and `Y` denote nonlinearity and pooling type, respectively (Section 5.2). The organization of panels is the same as that in Figure 6, except that only results for Monkey A are shown (see Figure 9 for Monkey B). Rows show different models, whose performance metrics (mean 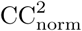) are decomposed into components of neuron subclasses, denoted by different colors (legend on the right). For each model in some panel, the length of each colored bar is equal to the average performance over that neuron subclass multiplied by the percentage of neurons in that subclass, and the length of all bars concatenated is equal to the average performance over all neurons. The baseline model has its name in bold, and “all” models in italics. d,e Neuron-by-neuron comparison of the a CNN variant with thresholding nonlinearity (`HS_max`) vs. one without (`S_max`) for OT neurons, all stimuli (d) and HO neurons, all stimuli (e). For d, e, and f, performance metrics (mean 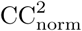) are shown at corners, Pearson correlation coefficients between models are shown at the top left, and regression lines for different neuron subclasses (colored solid) together with the regression line over all neurons (black dashed) are shown at the bottom right (scaled and shifted to the corner for clarity; otherwise these regression lines will clutter the dots that represent individual neurons). f Comparison of two thresholding nonlinearities, for HO neurons, all stimuli. Results with max pooling are shown, and average pooling gave similar results.

#### Thresholding nonlinearities outperformed non-thresholding ones

Compared to GLMs we explored in this work, one nonlinear structural component unique to CNNs is ReLU, a thresholding nonlinearity. To understand the usefulness of thresholding nonlinearities in general, we compared four CNN variants with thresholding nonlinearities (`R_max`, `R_avg`, `HS_max, HS_avg`) with four without (`A_max, A_avg, S_max, S_avg`) and found that thresholding (**R**, **HS**) in general helped. This can be seen at two levels. At the level of individual architectures, those with thresholding generally performed better than those without (d, e, and rows 5-8 from the top vs. 1-4 in a-c of Figures 8 and 9). At the level of model categories, we combined all four thresholding models into one “all” model (`T_all`) and all four non-thresholding ones as well (`NT_all`), using the same method as we constructed “all” models in Figure 6; we found that thresholding helped as well. Our results suggest that the recorded V1 neurons actually take some thresholded versions of the raw input stimuli as their own inputs. There are at least two ways to implement this input thresholding. First, neurons may have some other upstream neurons as their inputs, each upstream neuron with its own thresholding nonlinearity as modeled in McFarland et al. (2013); Vintch et al. (2015). Second, the thresholding may happen at the dendritic tree level, as suggested by Gollisch and Meister (2010).

**Fig. 9.**
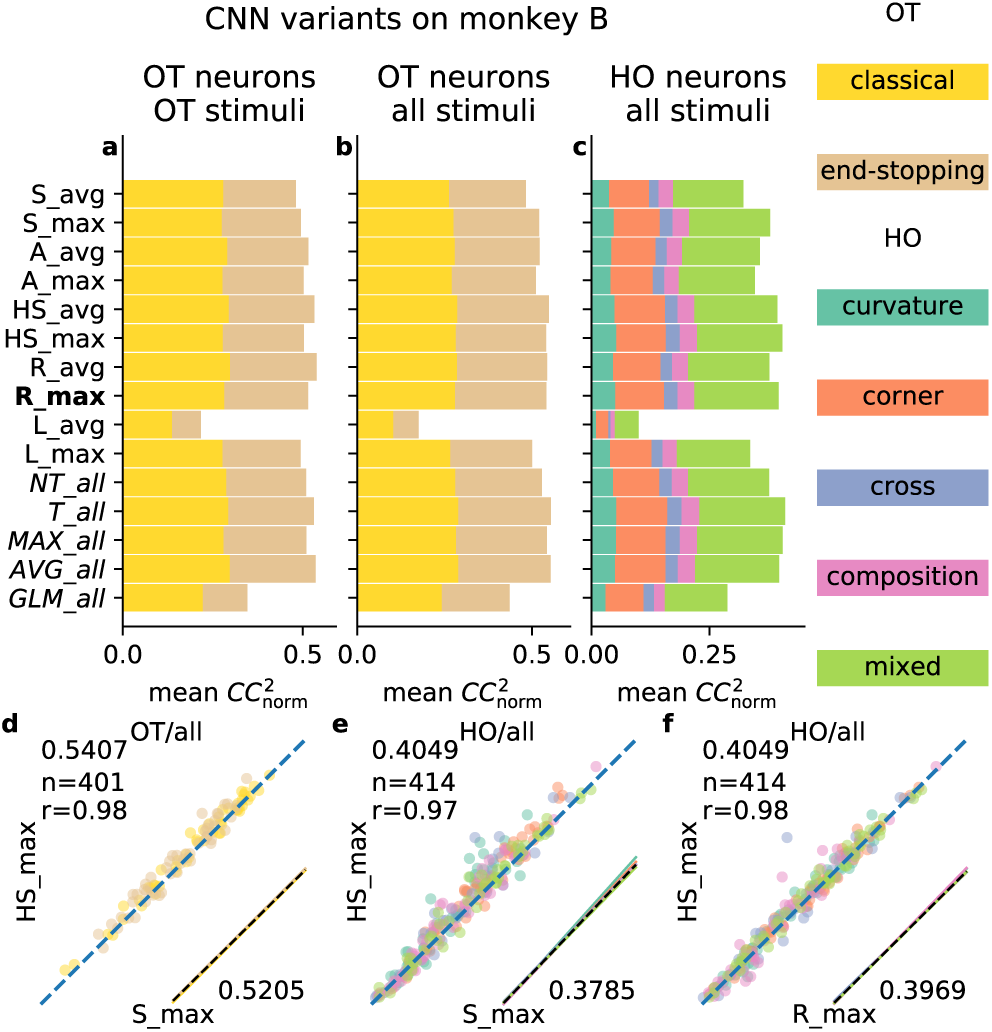
Detailed comparison of CNN variants, monkey B. Check Figure 8.

#### Thresholding nonlinearities performed similarly

While the two thresholding nonlinearities (`R` and `HS`) showed better performance overall, we did not see much difference between the two (`f`, and `HS_max` vs. `R_max`, `HS_avg` vs. `R_avg` in a-c of Figures 8 and 9). This observation was consistent with Heeger (1992), where the author claimed that these two types of thresholding nonlinearities are both consistent with physiological data and the brain might be using one as an approximation to implement the other.

#### No consistently better pooling type, but max pooling was more powerful in isolation

While thresholding nonlinearities showed better performance consistently than non-thresholding ones as shown above, the results were mixed for two pooling schemes and depended on non-linearities, combinations of neurons and stimuli, and monkeys (rows 1-8 from the top, as well as `MAX_all` vs. `AVG_all` that were constructed like `T_all` and `NT_all` above, in a-c of Figures 8 and 9). We suspect such mixed results were due to the complicated interaction between nonlinearity and pooling. In other words, the contributions of nonlinearity and pooling to model performance do not add linearly. Still, we think max pooling is a powerful computational component per se for modeling neural responses, as max pooling alone without any nonlinearity performed comparably with many other models with pooling and nonlinearity (`L_max` vs. others in a-c of Figures 8 and 9).

#### High correlation between per-neuron and average model performance

Figures 8 and 9 show that different models performed differently. We found that the performance increase/decrease of one model over another one seemed to be *universal*, rather than class- or neuron-specific. We can see this universality from several aspects when two models are compared neuron by neuron (d-f of Figures 8 and 9). First, there was a high correlation between the performance metrics of individual neurons (high Pearson correlation coefficients *r*). Second, we performed linear regression on each neuron subclass as well as on all neurons (colored solid lines and black dashed line in the lower right corner of each panel), and found all regression lines were very close.

### 5.3 Convolution was more effective than diverse filters

Apart from thresholding nonlinearity and max pooling explored in Section 5.2, CNN models have another unique structural component compared to other models in our study—shared weights among hidden units via convolution—as shown in Table 1. In contrast, other models with multiple hidden units (when these models are considered as MLPs; see Section 3.4) often have hidden units with independent and diverse weights without sharing (“independent” in Table 1). In this section, we explore the relative merits of these strategies for relating weights of different hidden units—shared weights via convolution vs. independent weights—in terms of model performance, not only for the CNN, but also for other model classes. The results are shown in Figure 10, with similar layout to Figures 8 and 9. We have the following observations (letters in the parentheses denote the panels used for highlighting).

– Multiple diverse filters alone did not help much (d vs. e).
– Convolution helped achieve better performance with the same number of parameters (f).

**Fig. 10.**
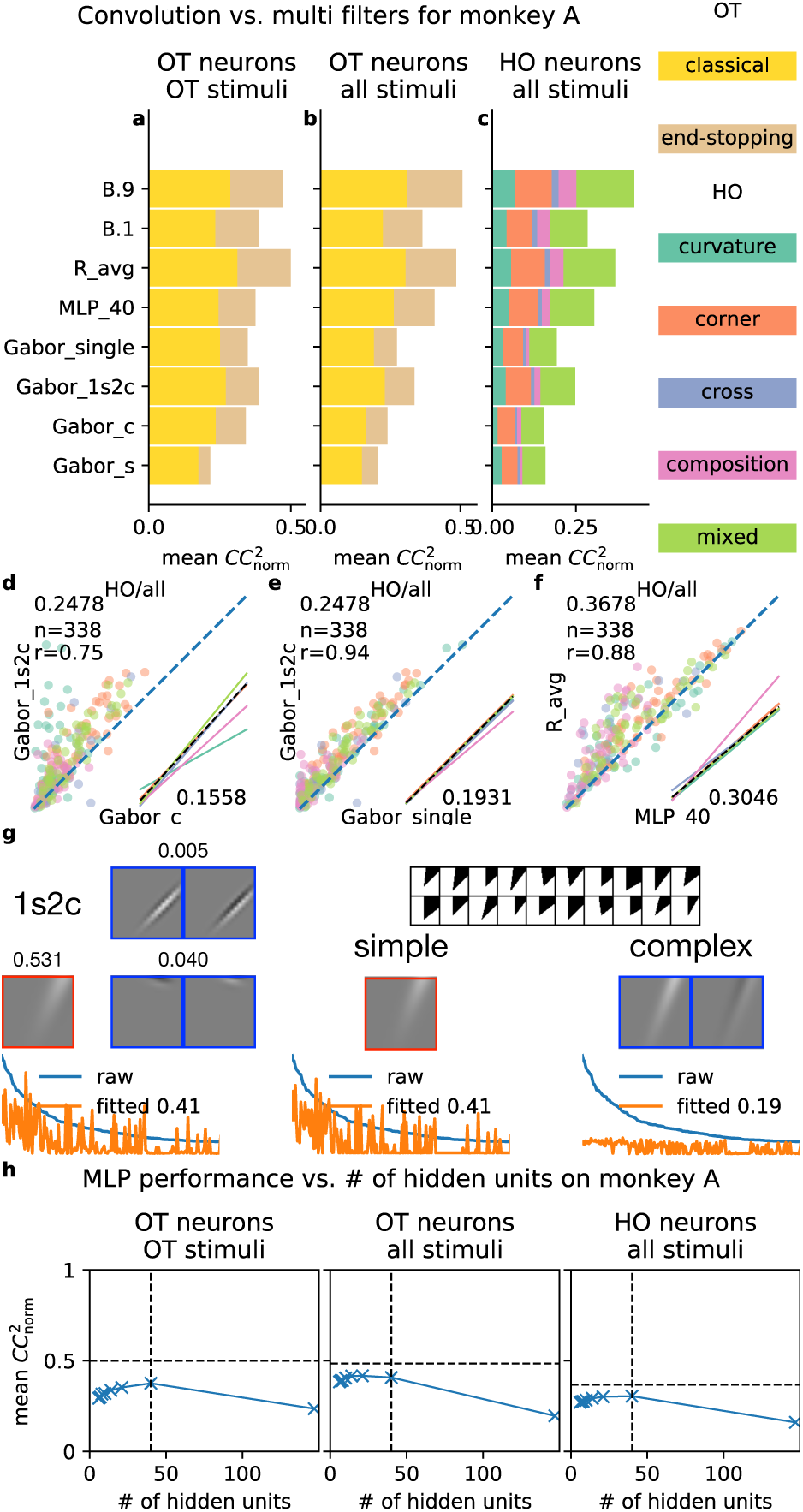
Convolution seemed more important than diverse filters. **a-f** Comparison of single-vs. multi-component Gabor models (highlighted in d,e), comparison of single-vs. multi-channel CNN models, and comparison of models with and without convolution (highlighted in **f**). See Section 5.3 for details. These panels have similar formats to those in Figure 8. **g** Learned single- (`simple` and `complex`) and multi-component (1s2c) Gabor models fitted to a particular neuron’s data. This neuron was tuned to corners as shown in the top right part of the panel. For the three models (left, middle, right), we show the learned filters (top) and fitting results (bottom). Simple cell components are shown with red borders, and complex cell components are shown with blue borders. For the multi-component model, we also show the weights of different com-ponents at the top of filters. In this case, the multi-component model was dominated by its simple component with weight 0.531, which was orders of magnitude larger than the weights of its complex components. **h** Performance vs. number of hidden units for MLP models. Vertical dashed lines denote the MLP model (`MLP_40`) in panels **a-c,f**, and horizontal dashed lines show performance metrics of the CNN `R_avg`. Only results for monkey A are shown and monkey B gave similar results.

#### Multiple diverse filters alone did not help much

To examine the impact of having multiple filters with diverse shapes, we explored two classes of models: Gabor models and CNN models. For Gabor models, we examined three single-filter variants—simple cell model (`Gabor_s`), complex cell model (`Gabor_c`), and the “single-mponent” Gabor model (`Gabor_single`) constructed from simple and complex cell models similarly to “all” models in Figure 6—and one multi-filter variant—one simple two complex (`Gabor_1s2c`; other multi-filter models performed worse as shown in Figure 6). For CNN models, we varied the number of channels of the baseline CNN B.9 from 1 (`B.1`) through 18 (`B.18`).

While the multi-filter Gabor model outperformed both simple and complex cell models by a large margin (a-c,d of Figure 10), we found that the single-component model (`Gabor_single`), which takes the better one of simple cell and complex cell models for each neuron, worked almost as well as the multi-filter one (a-c,e of Figure 10). While there was still some performance gap between `Gabor_single` and the `Gabor_1s2c`, the gap was relatively small and there was strong correlation between the two models in terms of per-neuron performance (Figure 10e). For each neuron, we further compared the learned filters of simple, complex, and multifilter Gabor models, and found that in some extreme cases, the learned multi-filter Gabor model was degenerate in the sense that it had its simple component dominate its complex components or vice versa (Figure 10g; check the caption).

The results for one-channel CNN and the baseline 9-channel CNN are shown in the top two rows of Figure 10a-c, and we found that the performance increase (around 20 % to 50 %) was not proportional to the increase in the number of parameters (around 800 %, or 99 vs. 883 parameters). See Figure 6 and Supplementary Materials for more results on the model performance of CNN as we change the number of channels.

#### Convolution helped achieve better performance with the same number of parameters

As shown in the previous part, having multiple independent filters of diverse shapes was not effective for increasing performance relative to the increase in model size it involved. However, we found that convolution was much more effective, achieving better model performance without increasing the number of parameters. To illustrate this, we compared the baseline CNN’s average pooling (R_avg)variant, which linearly combines ReLU units, with a multilayer perceptron consisting of one hidden layer of 40 ReLU units (MLP_40). To make the two models match in the number of parameters, we performed principal component analysis to reduce the input dimensionality for the MLP to 20; therefore the MLP has 40 × (20 + 1) + 40 + 1 = 881 parameters, roughly matching the CNN (883 parameters). The CNN outperformed the MLP by a relatively large margin (a-c,f of Figure 10). We also explored the trade-off between input dimensionality and number of hidden units for MLP, with the number of parameters roughly fixed (Figure 10h); given roughly the same number of parameters, the CNN, which has convolution, consistently outperformed MLPs of various configurations.

One may argue that the (average) pooling, which was difficult to avoid in our experiments as CNNs would otherwise have too many parameters, helped model performance as well; while such interpretation is possible, it is also helpful to simply consider convolution and pooling collectively as a modeling prior that helps neural response prediction with limited number of parameters and training data. The effectiveness of convolution and pooling could also be due to eye movements during neural data recording; as shown in our previous work (Tang et al., 2018), the eye movement was in general very small (the standard deviation of the distribution of eye positions during stimulus presentation was in general less than 0.05° in visual angle, or 0.75px in the 20 px by 20 px input space of the CNN) for our data, and such interpretation was less likely.

### 5.4 Data-driven vs. pre-trained CNNs and the complexity of V1 neural code

The results are shown in Figure 11. When different CNN layers are compared (Figure 11g), overall layer `conv3_1` performed the best (`conv4_1` performed similarly but we prefer layers that are lower and thus easier to analyze). The result was largely consistent with that in Cadena et al. (2017); however, we also observed performance decreases in layers `conv3_2` through `pool3` which were not present in Cadena et al. (2017), and we will investigate this in the future. When the best VGG layer and our baseline CNN B.9 are compared, the VGG layer performed simiarly to the CNN (Figure 11a-c), and there was a relatively high correlation between the performance metrics of individual neurons for two models (Figure 11d-f). We have also tried other variants of VGG and they performed similarly to or worse than VGG19 (Supplementary Materials).

**Fig. 11.**
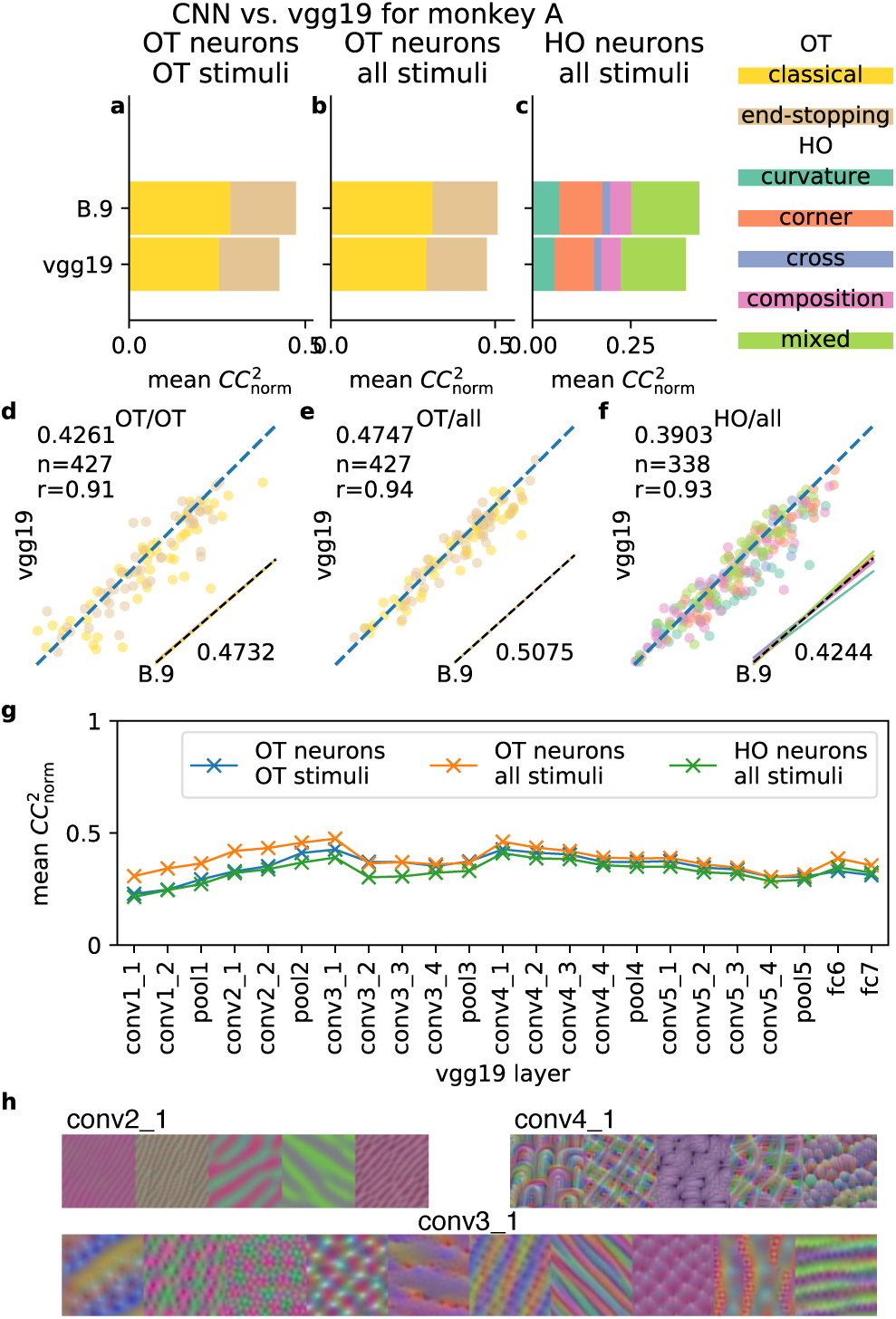
Transfer learning (goal-driven) approach for modeling V1 neurons using a pre-trained CNN (VGG19). a-f the best performing VGG19 layer (`conv3_l`) vs. the baseline CNN (B.9). These panels have similar formats to those in Figure 8. g Model performance across different VGG19 layers, for different combinations of neuron subsets and stimuli. Only results for monkey A are shown, and monkey B gave similar results (Supplementary Materials). h Visualization of some `conv2_1` (left), `conv3_1` (bottom), and `conv4_1` (right) units by activation maximization (Kotikalapudi, 2017; Olah et al., 2017). Each unit was visualized by the image (cropped to 50px by 50px) that maximizes the unit’s activation.

While our results show that pre-trained CNNs were on par with CNNs trained from stratch, it is possible that pre-trained CNNs would perform much better if they were trained on artificial stimuli as well due to the large difference between the image set used to train VGG19 (natural images) and from our artificial stimulus set. However, such possibility might be not very likely: (1) in our preliminary efforts to apply the state-of-the-art 3-layer CNN architecture in Cadena et al. (2017) to model our V1 neurons all together (Supplementary Materials), we found that the 3-layer CNN, within the limit of our hyperparameter tuning, performed similarly to our baseline CNN; (2) Cadena et al. (2017) have already established (somewhat) that the 3-layer CNN architecture and pre-trained CNNs perform similarly when all are trained with stimuli of similar nature. On the other hand, we found it interesting that CNNs trained on natural images could be used to effectively predict neural responses on artificial stimuli.

We also visualized units across various layers of the VGG19 network by activation maximization (Olah et al., 2017) implemented in `keras-vis` (Kotikalapudi, 2017), and found that these units from layers that matched neural data well (`conv3_1` and `conv4_1`) are tuned to relatively complex image features rather than oriented edges (Figure 11h); the visualization results were consistent with our earlier work (Tang et al., 2018) on the complexity of V1 neural code.

## 6 Discussion

### 6.1 Key components for the success of CNN

In this study, we evaluated a variety of Gabor-based models, generalized linear models, and CNN models for modeling V1 neurons of awake macaque monkeys. These models can be considered as a continuum of regression models in statistics or system identification models in sensory neuroscience (Wu et al., 2006); specifically, they can be considered as one-hidden-layer neural networks with different structural components and different degrees of model flexibility (Section 3.4). This comparative study allows us to empirically identify some key components that are important for modeling V1 neurons, particularly those neurons with selectivity to higher-order features as identified in Tang et al. (2018).

In Section 5.2, we evaluated CNN models under different combinations of nonlinearity (ReLU, half-squaring, squaring, and linear) and pooling scheme (max and average), and we found that thresholding nonlinearity and max pooling, which are absent in the best-performing non-CNN models (GLMs) in this study (Table 1), were important for the CNN’s superior performance relative to other models. In particular, we found that models with thresholding nonlinearities (ReLU and half-squaring) consistently performed better than those without. Interestingly, thresholding nonlinearities such as ReLU and half-squaring are already in classical models of simple cells (Andrews and Pollen, 1979; Heeger, 1992), and pooling (average pooling or max-pooling) of simple cells responses are also in some models of complex cells (Riesenhuber and Poggio, 1999; Finn and Ferster, 2007). In fact, these models of simple and complex cells were the inspiration to the development of the convolutional neural network architecture (Fukushima, 1980) in the first place. The Gabor-based models did not perform well mostly because their filters were restricted to Gabor functions whereas filters in 0 and CNNs could take arbitrary shapes. The CNN provides an elegant way of integrating nonlinearities in models of simple and complex cells with more flexible filters in GLMs; in addition, the CNN allows the linear combination of multiple filters, and such linear combination increases model expressiveness.

When all the models are considered as one-hidden-layer neural networks (Section 3.4), there are two strategies for relating weights learned for different hidden units—shared and spatially shifted weights (convolution) and independently learned weights for different units (Table 1, “constraints among units”). We evaluated the relative merits of these two strategies in Section 5.3 and found that convolution was more effective than having multiple independently learned units both for better performance and fewer model parameters. Our CNN models are similar to subunit models in the literature (Rust et al., 2005; Vintch et al., 2015; Hubel and Wiesel, 1962; McFarland et al., 2013), where V1 neurons take (thresholded) responses of upstream intracortical neurons (with approximately spatially shifted receptive fields) as their inputs and exhibit local spatial invariance, particularly for complex cells. Our CNN models are also consistent with previous V1 modeling work using spike-triggered methods (Rust et al., 2005; Park and Pillow, 2011) where the subspace spanned by recovered subunits can be approximated by the subspace spanned by one single set of spatially shifted subunits with shared weights. Despite the similarity between our CNN models and subunit models in the literature, our systematic and comprehensive exploration of the contribution of various structural components in the CNN helps to illuminate which nonlinear components are more important to the subunit models (Section 5.2) and what strategies for relating subunits were more effective (Section 5.3).

Overall, we believe this study is the first one that systematically evaluates the relative merits of different CNN components in the context of modeling V1 neurons. We demonstrated that key components of the CNN (convolution, thresholding nonlinearity, and pooling) contributed to its superior performance in explaining V1 responses. Our results suggests that that there is a high degree of correspondence between the CNN and biological reality.

### 6.2 Complexity of V1 neural code

Using our neural dataset (Section 2), we have found earlier (Tang et al., 2018) that a large proportion of V1 neurons in superficial layers are selective to higher-order complex features rather than simple oriented edges. We classified these neurons selective to higher-order complex features as “higher-order” (HO) neurons and others as “orientation-tuned” (OT) neurons (Section 2), some of which also exhibited complex feature selectivities due to the strictness of our classification criterion (Tang et al., 2018). In this study, we showed that the performance gap between CNN models and non-CNN models (Gabor-based models and GLMs) was more pronounced in HO neurons than in OT neurons, and we took this as an additional evidence supporting the complexity of V1 neural code in HO neurons.

In addition, by fitting intermediate features from a pre-trained, goal-driven CNN (VGG19) to our neural data, we found that a relatively high layer (`conv3_1`), which encodes relatively complex image features (Figure 11h), explained our neural data the best among all the VGG19 layers. This finding further reinforced the claim that V1 neurons in superficial layers might have a great degree of complex selectivity and was largely consistent with Cadena et al. (2017) where the same layer (conv3_1) in VGG19 provides the best model of V1 responses to natural images in their study. Furthermore, the fact that such pre-trained, goal-driven neural networks performed well for explaining neural responses both in V1 (our study and Cadena et al. (2017)) and in IT (Yamins and DiCarlo, 2016; Yamins et al., 2013; Kriegeskorte, 2015) is another piece of evidence that there is a high degree of correspondence between the CNN and biological reality.

### 6.3 Limitations and varieties of the CNN

While CNN models, especially those goal-driven ones pre-trained on computer vision tasks, performed very well in our study and some other studies (Cadena et al., 2017) for V1 neuron modeling, we should point out that even the best-performing CNN in our study only explained about 50 % of the explainable variance in our neural data, consistent with Cadena et al. (2017). The failure of CNN models for explaining the other half of the variance in V1 data can be due to a number of reasons. First, V1 neurons are subject to network interaction and their neural responses are known to be mediated by strong long-range contextual modulation. Second, it is possible that there are some basic structural components missing in the current deep CNN methodology for fully capturing V1 neural code.

An important design issue with CNN modeling is the depth of the network. Here, we used a very basic one-convolutional-layer CNN because we can then formulate all models as one-hidden-layer neural networks (Section 3.4) and directly evaluate the relative contributions of different model components (Sections 5.2 and 5.3). We found that, at least for our data, adding an additional convolutional layer did not produce significant performance benefit (Supplementary Materials), and the performance difference between our baseline one-convolutional-layer CNN and the pre-trained CNN was relatively small (Figure 11). Our findings suggest that the true nature of V1 neurons does not need to be modeled by a very deep network. However, other studies typically use deeper CNNs. McIntosh et al. (2017) used a 2-convolutional-layer CNN with to model retinal ganglion cells; Kindel et al. (2017) and Cadena et al. (2017) used 2- and 3-convolutional-layer CNNs to model V1 neurons respectively. It is possible that 2 or 3-layer networks are needed for modeling V1 neural responses to natural images that are richer than our artificial stimuli; in addition, higher layers in those models might be functionally equivalent to the max pooling layer in our CNN, as those multi-layer CNNs typically do not use pooling. Given there are multiple layers of neurons on the pathway between the photoreceptors in the retina and superficial layer cells in V1, biologically speaking, a much deeper CNN should provide a more accurate model in a more general setting.

Most CNN-based work models all the neurons in a data set with a single network, with shared parameters in lower layers and separate sets of parameters for different neurons in higher layers (Kindel et al., 2017; McIntosh et al., 2017; Klindt et al., 2017; Cadena et al., 2017). Instead, we model each neuron separately to allow a more fair comparison between CNN models and other models (GLMs, etc.) that typically model neurons separately without parameter sharing, and a fair comparison allows us to understand the CNN’s success compared to other models more conveniently. We also tried modeling all neurons using a single CNN (Supplementary Materials) with an architecture similar to that in Cadena et al. (2017); to our surprise, large single CNNs that model all neurons together performed similarly to our baseline CNNs that model each neuron separately, given roughly the same number of parameters; for example, a 3-layer single CNN with around 300k parameters trained on some (around 350) HO neurons performed similarly to separately trained CNNs, which together take around 300k parameters (883 parameters per model) as well. More investigation (better hyperparameter tuning, better network architecture, etc.) is needed to improve the performance of modeling using a single CNN on our V1 data.

## Acknowledgments

We thank Rob Kass, Dave Touretzky, and the reviewers for careful comments on the manuscript. We thank Wenbiao Gan for the early provision of AAV-GCaMP5; and to Peking University Laboratory Animal Center for excellent animal care. We acknowledge the Janelia Farm program for providing the GCaMP5-G construct, specifically Loren L. Looger, Jasper Akerboom, Douglas S. Kim, and the Genetically Encoded Calcium Indicator (GECI) project at Janelia Farm Research Campus Howard Hughes Medical Institute. This work was supported by the National Natural Science Foundation of China No. 31730109, National Natural Science Foundation of China Outstanding Young Researcher Award 30525016, a project 985 grant of Peking University, Beijing Municipal Commission of Science and Technology under contract No. Z151100000915070, NIH 1R01EY022247 and NSF CISE 1320651 and IARPA D16PC00007 of the U.S.A.

## Footnotes

1 the images were rescaled to 2=3 of their original sizes; we used this scale because in another study (Zhang et al., 2016) we found that this scale gave the highest representational similarity (Kriegeskorte et al., 2008) between the CNN and neural data among all scales explored; we also tried using raw images without rescaling in the current study and got worse results.

2 in theory we should exclude these neurons for model evaluation, we did not do it as doing it or not has negligible effects with hundreds of neurons in our data set.

3 In practice, we performed PCA only on the pure quadratic terms to reduce their dimensionalities to 432 and concatenated the PCAed 432-dimensional pure quadratic terms with the 400-dimensional linear terms to generate the final 882dimensional input vectors; such method would guarantee that the information from linear terms, which are heavily used in most V1 models, is preserved. We also tried performing PCA on both linear and pure quadratic terms together and two methods made little difference in our experiments.

## References

Adelson, E. H. and Bergen, J. R. (1985). Spatiotemporal energy models for the perception of motion. J. Opt. Soc. Am., 2(2):284–299.

Andrews, B. W. and Pollen, D. A. (1979). Relationship between spatial frequency selectivity and receptive field profile of simple cells. The Journal of Physiology, 287(1):163–176.

Bishop, C. M. (2006). Machine learning and pattern recognition. Information Science and Statistics. Springer.

Cadena, S. A., Denfield, G. H., Walker, E. Y., Gatys, L. A., Tolias, A. S., Bethge, M., and Ecker, A. S. (2017). Deep convolutional models improve predictions of macaque v1 responses to natural images. bioRxiv.

Carandini, M., Demb, J. B., Mante, V., Tolhurst, D. J., Dan, Y., Olshausen, B. A., Gallant, J. L., and Rust, N. C. (2005). Do We Know What the Early Visual System Does? Journal of Neuroscience, 25(46):10577–10597.

Coen-Cagli, R., Kohn, A., and Schwartz, O. (2015). Flexible gating of contextual influences in natural vision. Nature Neuroscience, 18:1648 EP –.

Daugman, J. G. (1985). Uncertainty relation for resolution in space, spatial frequency, and orientation optimized by two-dimensional visual cortical filters. J. Opt. Soc. Am. A, 2(7):1160–1169.

David, S. V. and Gallant, J. L. (2005). Predicting neuronal responses during natural vision. Network: Computation in Neural Systems, 16(2–3):239–260.

Dayan, P. and Abbott, L. F. (2001). Theoretical Neuroscience. Computational and Mathematical Modeling of Neural Systems.

Finn, I. M. and Ferster, D. (2007). Computational diversity in complex cells of cat primary visual cortex. Journal of Neuroscience, 27(36):9638–9648.

Friedman, J., Hastie, T., and Tibshirani, R. (2010). Regularization paths for generalized linear models via coordinate descent. Journal of Statistical Software, Articles, 33(1):1–22.

Fukushima, K. (1980). Neocognitron: A self-organizing neural network model for a mechanism of pattern recognition unaffected by shift in position. Biological Cybernetics, 36(4):193–202.

Gollisch, T. and Meister, M. (2010). Eye smarter than scientists believed: Neural computations in circuits of the retina. Neuron, 65(2):150–164.

Goodfellow, I. J., Bengio, Y., and Courville, A. (2016). Deep Learning. MIT Press.

Heeger, D. J. (1992). Half-squaring in responses of cat striate cells. Visual Neuroscience, 9(5):427443.

Hegdé, J. and Van Essen, D. C. (2007). A comparative study of shape representation in macaque visual areas v2 and v4. Cerebral Cortex, 17(5):1100–1116.

Hsu, A., Borst, A., and Theunissen, F. (2004). Quantifying variability in neural responses and its application for the validation of model predictions. Network: Computation in Neural Systems, 15(2):91–109.

Hubel, D. H. and Wiesel, T. N. (1959). Receptive fields of single neurones in the cat’s striate cortex. The Journal of Physiology, 148(3):574–591.

Hubel, D. H. and Wiesel, T. N. (1962). Receptive fields, binocular interaction and functional architecture in the cat’s visual cortex. The Journal of Physiology, 160(1):106–154.

Hubel, D. H. and Wiesel, T. N. (1968). Receptive fields and functional architecture of monkey striate cortex. The Journal of Physiology, 195(1):215–243.

Jones, J. P. and Palmer, L. A. (1987a). An evaluation of the two-dimensional Gabor filter model of simple receptive fields in cat striate cortex. Journal of Neurophysiology, 58(6):1233–1258.

Jones, J. P. and Palmer, L. A. (1987b). The two-dimensional spatial structure of simple receptive fields in cat striate cortex. Journal of Neurophysiology, 58(6):1187–1211.

Kelly, R. C., Smith, M. A., Kass, R. E., and Lee, T. S. (2010). Accounting for network effects in neuronal responses using L1 regularized point process models. In Lafferty, J. D., Williams, C. K. I., Shawe-Taylor, J., Zemel, R. S., and Culotta, A., editors, Advances in Neural Information Processing Systems 23: 24th Annual Conference on Neural Information Processing Systems 2010. *Proceedings of a meeting held 6-9* December 2010, Vancouver, British Columbia, Canada., pages 1099–1107. Curran Associates, Inc.

Kindel, W. F., Christensen, E. D., and Zylberberg, J. (2017). Using deep learning to reveal the neural code for images in primary visual cortex. ArXiv e-prints, q-bio.NC.

Kingma, D. P. and Ba, J. (2014). Adam: A method for stochastic optimization. CoRR, abs/1412.6980.

Klindt, D., Ecker, A. S., Euler, T., and Bethge, M. (2017). Neural system identification for large populations separating ‘what’ and ‘where’. In Guyon, I., von Luxburg, U., Bengio, S., Wallach, H. M., Fergus, R., Vishwanathan, S. V. N., and Garnett, R., editors, Advances in Neural Information Processing Systems 30: Annual Conference on Neural Information Processing Systems 2017, 4-9 December 2017, Long Beach, CA, USA, pages 3509–3519.

Köster, U. and Olshausen, B. (2013). Testing our conceptual understanding of V1 function. ArXiv e-prints, q-bio.NC.

Kotikalapudi, R. (2017). keras-vis: Keras visualization toolkit. https://github.com/raghakot/keras-vis.

Kriegeskorte, N. (2015). Deep Neural Networks: A New Framework for Modeling Biological Vision and Brain Information Processing. Annual Review of Vision Science, 1(1):417–446.

Kriegeskorte, N., Mur, M., and Bandettini, P. (2008). Representational similarity analysis connecting the branches of systems neuroscience. Frontiers in Systems Neuroscience, 2:4.

Krizhevsky, A., Sutskever, I., and Hinton, G. E. (2012). Imagenet classification with deep convolutional neural networks. In Bartlett, P. L., Pereira, F. C. N., Burges, C. J. C., Bottou, L., and Weinberger, K. Q., editors, Advances in Neural Information Processing Systems 25: 26th Annual Conference on Neural Information Processing Systems 2012. *Proceedings of a meeting held* December 3-6, 2012, Lake Tahoe, Nevada, United States., pages 1106–1114.

Li, M., Liu, F., Jiang, H., Lee, T. S., and Tang, S. (2017). Long-term two-photon imaging in awake macaque monkey. Neuron, 93(5):1049–1057.e3.

McCullagh, P. and Nelder, J. (1989). Generalized Linear Models, Second Edition. Chapman & Hall/CRC Monographs on Statistics & Applied Probability. Taylor & Francis.

McFarland, J. M., Cui, Y., and Butts, D. A. (2013). Inferring Nonlinear Neuronal Computation Based on Physiologically Plausible Inputs. PLoS computational biology, 9(7):e1003143.

McIntosh, L. T., Maheswaranathan, N., Nayebi, A., Ganguli, S., and Baccus, S. A. (2017). Deep Learning Models of the Retinal Response to Natural Scenes. ArXiv e-prints, q-bio.NC.

Olah, C., Mordvintsev, A., and Schubert, L. (2017). Feature visualization. Distill. https://distill.pub/2017/feature-visualization.

Paninski, L. (2004). Maximum likelihood estimation of cascade point-process neural encoding models. Network: Computation in Neural Systems, 15(4):243–262.

Park, I. M., Archer, E., Priebe, N., and Pillow, J. W. (2013). Spectral methods for neural characterization using generalized quadratic models. In Burges, C. J. C., Bottou, L., Ghahramani, Z., and Weinberger, K. Q., editors, Advances in Neural Information Processing Systems 26: 27th Annual Conference on Neural Information Processing Systems 2013. *Proceedings of a meeting held* December 5-8, 2013, Lake Tahoe, Nevada, United States., pages 2454–2462.

Park, I. M. and Pillow, J. W. (2011). Bayesian Spike-Triggered Covariance Analysis. In Taylor, J. S., Zemel, R. S., Bartlett, P. L., Pereira, F. C. N., and Weinberger, K. Q., editors, Advances in Neural Information Processing Systems 24: 25th Annual Conference on Neural Information Processing Systems 2011. *Proceedings of a meeting held 12-14* December 2011, Granada, Spain., pages 1692–1700.

Paszke, A., Gross, S., Chintala, S., Chanan, G., Yang, E., DeVito, Z., Lin, Z., Desmaison, A., Antiga, L., and Lerer, A. (2017). Automatic differentiation in pytorch.

Pillow, J. W., Shlens, J., Paninski, L., Sher, A., Litke, A. M., Chichilnisky, E. J., and Simoncelli, E. P. (2008). Spatio-temporal correlations and visual signalling in a complete neuronal population. Nature, 454:995 EP –.

Prenger, R., Wu, M. C. K., David, S. V., and Gallant, J. L. (2004). Nonlinear V1 responses to natural scenes revealed by neural network analysis. Neural Networks, 17(5–6):663–679.

Riesenhuber, M. and Poggio, T. (1999). Hierarchical models of object recognition in cortex. Nature Neuroscience, 2:1019 EP –.

Rowekamp, R. J. and Sharpee, T. O. (2017). Cross-orientation suppression in visual area V2. Nature Communications, 8:15739.

Rust, N. C., Schwartz, O., Movshon, J. A., and Simoncelli, E. P. (2005). Spatiotemporal Elements of Macaque V1 Receptive Fields. Neuron, 46(6):945–956.

Schoppe, O., Harper, N. S., Willmore, B. D. B., King, A. J., and Schnupp, J. W. H. (2016). Measuring the Performance of Neural Models. Frontiers in Computational Neuroscience, 10:1929.

Simonyan, K. and Zisserman, A. (2014). Very Deep Convolutional Networks for Large-Scale Image Recognition. ArXiv e-prints, cs.CV.

Tang, S., Lee, T. S., Li, M., Zhang, Y., Xu, Y., Liu, F., Teo, B., and Jiang, H. (2018). Complex Pattern Selectivity in Macaque Primary Visual Cortex Revealed by Large-Scale Two-Photon Imaging. Current Biology, 28(1):38–48.e3.

Theunissen, F. E., David, S. V., Singh, N. C., Hsu, A., Vinje, W. E., and Gallant, J. L. (2001). Estimating spatio-temporal receptive fields of auditory and visual neurons from their responses to natural stimuli. Network: Computation in Neural Systems, 12(3):289–316.

Touryan, J., Felsen, G., and Dan, Y. (2005). Spatial Structure of Complex Cell Receptive Fields Measured with Natural Images. Neuron, 45(5):781–791.

Victor, J. D., Mechler, F., Repucci, M. A., Purpura, K. P., and Sharpee, T. (2006). Responses of v1 neurons to two-dimensional hermite functions. Journal of Neurophysiology, 95(1):379–400.

Vintch, B., Movshon, J. A., and Simoncelli, E. P. (2015). A convolutional subunit model for neuronal responses in macaque v1. Journal of Neuroscience, 35(44):14829–14841.

Wu, M. C. K., David, S. V., and Gallant, J. L. (2006). Complete Functional Characterization of Sensory Neurons by System Identification. Annual Review of Neuroscience, 29(1):477–505.

Yamins, D., Hong, H., Cadieu, C. F., and DiCarlo, J. J. (2013). Hierarchical Modular Optimization of Convolutional Networks Achieves Representations Similar to Macaque IT and Human Ventral Stream. In Burges, C. J. C., Bottou, L., Ghahramani, Z., and Weinberger, K. Q., editors, Advances in Neural Information Processing Systems 26: 27th Annual Conference on Neural Information Processing Systems 2013. *Proceedings of a meeting held* December 5-8, 2013, Lake Tahoe, Nevada, United States., pages 3093–3101.

Yamins, D. L. K. and DiCarlo, J. J. (2016). Using goal-driven deep learning models to understand sensory cortex. Nature Neuroscience, 19(3):356–365.

Zhang, Y., Massot, C., Zhi, T., Papandreou, G., Yuille, A., and Lee, T. S. (2016). Understanding neural representations in early visual areas using convolutional neural networks. In Neuroscience (SfN).

